# Cell shape noise strength regulates the rate of shape change during EMT-associated cell spreading

**DOI:** 10.1101/2024.10.14.618199

**Authors:** Wolfram Pönisch, Iskra Yanakieva, Guillaume Salbreux, Ewa K. Paluch

## Abstract

Cellular shape is intimately linked to cell function and state, and transitions between cell states are tightly coupled to shape changes. Yet, shape has been largely overlooked in state transitions studies. Here, we combine morphometric analysis with theoretical modeling and molecular perturbations to interrogate cell shape dynamics during epithelial-to-mesenchymal transition (EMT). Using stochastic inference, we extract the morphogenetic landscape underlying EMT. We show that within this landscape, EMT-associated cell spreading reflects a transition between shape attractors. Strikingly, we observe a peak in cell shape noise strength concomitant with spreading, and show that higher shape noise accelerates transitions between shape attractors. Our morphometric analysis framework will be widely applicable to quantitative cell shape investigations in physiology and disease. Together, our results identify a key role for cellular stochasticity as a regulator of shape change rate, and highlight that shape dynamics yield rich phenotypic information that enhances our understanding of cellular states.

## INTRODUCTION

Most cell types in an organism display functionally adapted, well-defined shapes and shape deregulation is a harbinger of disease from developmental disorders to cancer^1–3^, a feature utilized in histological diagnostics. As such, precise control of cell shape is a key aspect of cell function, and morphological changes accompany most cellular state and fate transitions during embryonic development, tissue homeostasis and disease progression.

Cell state transitions have been extensively studied. However, most studies focus on underlying molecular changes and do not consider cell shape. As a result, cell state transitions are primarily delineated through molecular changes, at the level of biochemical signalling or gene and protein expression. In particular, large scale omics analyses combined with advanced bioinformatics pipelines have established highly quantitative frameworks for the definition of cell states based on differences in molecular signatures^4^. In this context, state transitions are often described using the metaphor of Waddington landscapes, where the continuum of molecular configurations is likened to a landscape with peaks and valleys. Valleys in such landscapes represent attractor states, describing stable or favorable molecular patterns towards which cells are drawn, and cell state transitions are described as transitions between valleys. Mathematically, this concept of state attractors has provided a powerful framework, yielding many insights into state determination and the mechanisms of cell state change at the molecular level^5–7^.

In contrast, changes in cell morphology associated with state transitions have received much less attention. This is partly because cell shape changes ultimately result from changes in cell mechanics and in forces exerted on the cell surface. While mechanical changes can be triggered by changes in signaling or gene expression, they are mediated by physical reorganization of cytoskeletal architecture, of the plasma membrane, and other mechanically active cellular structures^8–10^. Understanding how these mechanical properties relate to gene expression changes is challenging and remains poorly understood. As a result, morphological changes associated with cell state transitions have been largely overlooked.

Another obstacle in investigating morphological transitions is the technical challenge of quantifying cell shape, particularly in live cells. Historically, shape changes have mostly been monitored using hand-picked descriptors of interest, such as aspect ratio or projected area to describe cell spreading. While such measures can be informative^11,12^, they provide only limited information on overall cell morphology. In recent years, enhanced capabilities of imaging and segmentation have been enabling more systematic approaches to cell shape investigations based on morphometrics. In these approaches, cell shape is quantified through a high-dimensional set of features, either intuitive^13–18^ or using mathematical descriptors^19–26^. Using dimensionality reduction, shape features are then mapped into a low-dimensional morphospace, which quantitatively describes cell shapes. These new developments provide a mathematical framework for the quantitative investigation of cellular shape changes.

Recent studies have used morphospace analysis to investigate epithelial-to-mesenchymal transition (EMT)^27,28^, a phenotypic state change of key importance for development, wound healing and disease^29,30^. During EMT, cells transition from compact epithelial morphologies to spread cell shapes and increased cell motility (Figure 1A). These shape changes enable cells of epithelial origin to migrate away from their original position, a key function of EMT^30^. As such, the extent of cell shape change has been used as a single cell dynamic readout of EMT progression^31,27^. Furthermore, a combined analysis of 2D cell shape trajectories and intracellular distributions of mesenchymal markers suggests that EMT is driven by a transition between attractors in this composite feature space^27^. To what extent attractors can be identified when considering cell shape alone, and how cells dynamically transition between such attractors has, to our knowledge, not been investigated. As a result, the mechanisms controlling the dynamics of shape change in EMT remain poorly understood.

**Figure 1.**
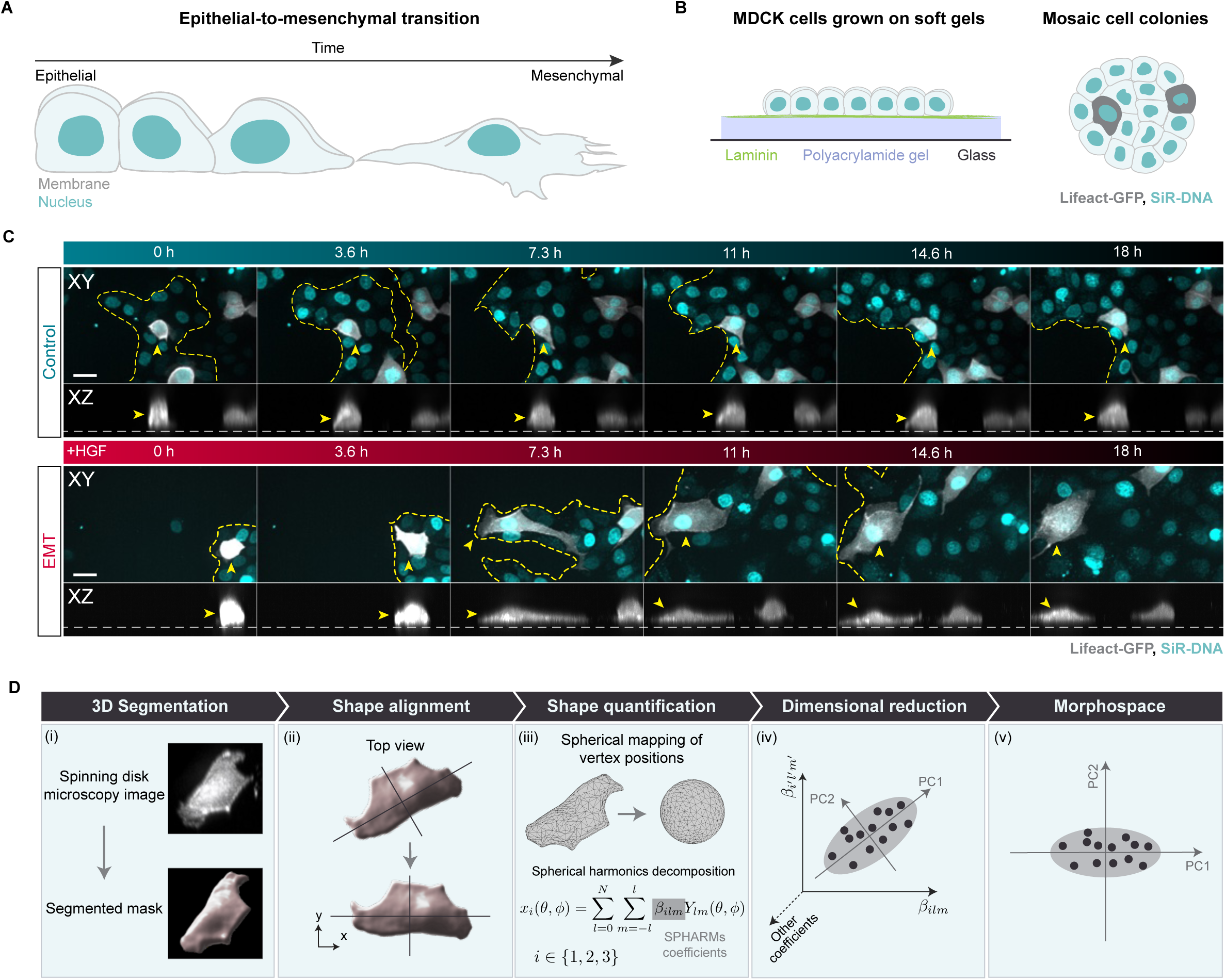
Morphometric analysis of cell shape during EMT. (A) Schematic of EMT-associated cell shape changes. (B) Schematic of the experimental approach. MDCK cells were cultured on soft polyacrylamide gels coated with laminin to promote cuboidal epithelial morphologies prior to EMT induction. Mosaic cell labeling was achieved by mixing MDCK cells stably expressing Lifeact-GFP (gray) with unlabeled MDCK cells (light blue). All nuclei were labelled with SiR-DNA (cyan). (C) Representative time series of control and EMT cells, imaged with spinning disk confocal microscope. XY – top view, XZ – side view in maximum intensity projections. Labels: Lifeact-GFP (F-actin, gray) and SiR-DNA labels (nuclei, cyan). Yellow dashed line: colony outline, arrowhead: representative Lifeact-GFP cell. Scale bars: 20 µm. (D) Morphometric analysis pipeline for quantitative cell shape description. (i) Segmentation of 3D cell shapes from experimental data. (ii) Alignment of cell shape so that the long axis of all cells is oriented along the *x*-axis. (iii) Decomposition of cell shape into SPHARMs coefficients *β*_*ilm*_. (iv) Dimensional reduction of high-dimensional data set of SPHARM coefficients *β*_*ilm*_using principal component analysis. (v) Low-dimensional mapping of cell shape in a morphospace. See also Methods for details on the analysis pipeline.

Here, we investigated 3D cell shape dynamics during EMT and asked whether the shape change can be described as a transition between shape attractors in a morphogenetic landscape. We used Madin-Darby canine kidney (MDCK) epithelial cells, widely used in studies of EMT^32,33^, and developed a morphospace analysis pipeline to investigate the stochastic dynamics of complex 3D cell shape change. We then used stochastic inference to extract the morphogenetic landscape and the strength of cell shape noise underlying morphospace shape trajectories. We found that EMT-associated cell spreading can be mathematically accounted for by a change in shape attractor. Strikingly, we also observed a strong increase in cell shape noise at the time of the shape transition. Theoretical analysis and perturbation experiments strongly suggest that cell shape noise accelerates cell spreading. Together, our study shows that EMT-associated cell spreading can be mathematically described as a transition between cell shape attractors, and that enhanced cell shape noise facilitates this transition, thus playing a key role in controlling cell shape change.

## RESULTS

### Generating a morphospace for the analysis of EMT-associated cell shape dynamics

We first developed a morphometric analysis pipeline to investigate 3D cell shape changes displayed by MDCK cells during EMT (Figures 1 and S1). We cultured cells on soft polyacrylamide hydrogels^34^ to promote cuboidal cell morphologies in the epithelial state^35^ (Figure 1B), and induced EMT by adding Hepatocyte Growth Factor (HGF)^33^. To facilitate cell shape segmentation, we used a mosaic labeling approach, where only a subset of cells expresses the F-actin marker Lifeact-GFP (Figures 1B and 1C). We acquired 3D image stacks for up to 22 h after EMT induction and observed that while control cells remained cuboidal throughout the imaging window, cells undergoing EMT displayed dramatic shape changes of the whole cell body as they spread on the substrate (Figure 1C, Movie S1).

We used morphospace representation to quantitatively characterize 3D shape dynamics during EMT (Figure 1D; Methods). Cell segmentation yielded timecourses of 3D triangulated meshes (Figure 1D(i)). We first aligned the long cell axes by rotating cell meshes in the plane of the substrate, so that a cell’s orientation did not impact its position in morphospace (Figure 1D(ii)). We then mapped the cell surface mesh points onto a sphere and performed a spherical harmonics (SPHARMs) decomposition to extract a list of SPHARMs coefficients that represent the cell shape, providing a robust, automated quantification of cell morphology (Figures 1D(iii) and S1; Methods). We found that SPHARMs functions up to a degree of 23, corresponding to 1728 coefficients, were sufficient to reconstruct cell shapes in our dataset with a high degree of accuracy (Figures S1A-S1D). To facilitate the visualization and interpretation of this high-dimensional dataset, we performed dimensionality reduction using principal component analysis (PCA; Figure 1D(iv); Methods), enabling the representation of cell shapes as points in a low-dimensional morphospace (Figure 1D(v)). We applied PCA on a dataset of cells undergoing EMT at 0 h and 18 h after EMT induction (0 h and 18 h EMT cells thereafter), as these time points reflect the extreme stages of the shape trajectories, corresponding to epithelial and mesenchymal morphologies, respectively (Figure 1C). This determined the principal components (PCs) of cell shape variation during EMT, which we used as axes of a morphospace to mathematically represent and analyze cell shapes in the rest of the study.

### Morphometric analysis of EMT-associated cell shape change

We then characterized EMT cell shape changes using our morphospace representation. We first plotted control and EMT cell shapes at 0 h and 18 h in the two-dimensional PC1-PC2 morphospace (Figures 2A and S2A-S2D). We found that for control cells, the mean position and the PC1-PC2 cell shape distribution did not change considerably over time (Figure 2A and S2A). In contrast, while the cell shape distribution of EMT cells overlapped with that of control cells at 0 h (Figure S2C), 18 h EMT cells displayed a clear shift towards higher values of PC1 and a wider PC1-PC2 cell shape distribution compared to 0 h EMT cells (Figure 2A and S2D). Visual analysis indicates that an increase in PC1 reflects changes in multiple shape features associated with cell spreading, including an increase in projected area and a decrease in cell height, while an increase in PC2 reflects asymmetric cell elongation (Figures 2B, S2E, S2F, Movie S2, Movie S3).

**Figure 2.**
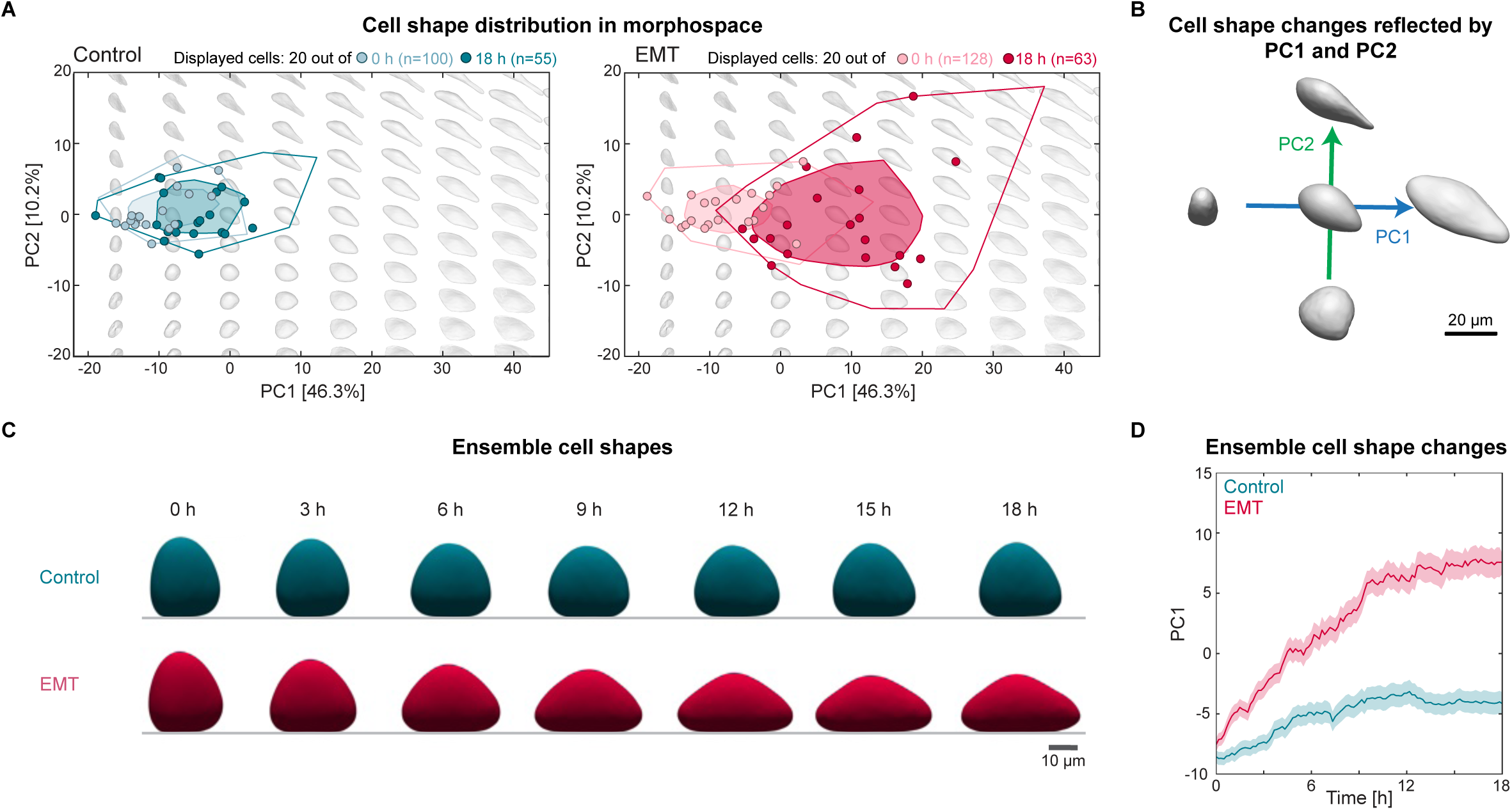
Ensemble cell shape changes during EMT. (A) Distribution of 0 h and 18 h control (N=6, n=100 and N=6, n=55) and EMT cells (N=10, n=128 and N=10, n=63) in the PC1-PC2 morphospace. For each condition, we show the morphospace positions of 20 randomly chosen cells (see Figures S2A and S2B for all datapoints). Background gray shapes: reconstructed theoretical cell shapes for different combinations of PC1 and PC2. Datapoints: individual cell shapes. Polygons: bagplots^64^, where the inner polygon (bag) is constructed based on the Tukey depth and contains at most 50% of the points and the outer polygon (fence) corresponds to the convex hull of all datapoints. (B) Reconstruction of cell shape changes reflected by changes in PC1 and PC2, see Methods for details of the reconstruction. (C) Mean shape dynamics of control and EMT cells. The mean shapes were generated by averaging the SPHARMs coefficients *β*_*ilm*_(*t*) of all cells at the given times and reconstructing the shape associated with the average SPHARM coefficient 〈*β*_*ilm*_(*t*)〉 (Methods). (D) Ensemble dynamics of PC1 of control (N=6, n=102) and EMT cells (N=12, n=145). Plots: mean ± SEM.

We then analyzed the temporal dynamics of the main PCs of shape variation. As a reference, we characterized ensemble cell shape dynamics by averaging the complete set of SPHARMs coefficients and reconstructed the resulting morphologies, yielding ensemble-averaged cell shapes over the imaging timecourse (Figure 2C). While the reconstructed average control cell did not display any major shape change, the average EMT cell exhibited spreading within 9 to 12 h after EMT induction. We next asked how average PC dynamics related to overall cell shape dynamics. We found that while the average PC1 increased weakly with time for control cells, average PC1 of EMT cells displayed a steep increase after EMT induction and saturated after 10-12 h (Figure 2D). In contrast, average PC2 dynamics initially displayed no difference between control and EMT cells, and showed a weak difference after 9 h (Figure S2G), while average PC3 displayed only very minor differences between EMT and control cells over the total timecourse (Figure S2H). Together with the morphospace analysis (Figure 2A), these observations suggest that PC1 captures most of the morphological changes associated with EMT.

We then quantitatively assessed how many PCs were required to account for EMT-associated cell shape changes. Cumulative contribution analysis showed that 22 PCs are needed to account for more than 90% of the total variance in our dataset of EMT cell shapes at 0 h and 18 h (Figure S1E). While part of this whole dataset variance reflects EMT-associated cell shape change, some of the variance corresponds to shape variability between cells at any given timepoint. To disentangle these contributions, we compared the standard deviations of each PC values for EMT cells at 0 h and 18 h (accounting for shape variability at 0 h and 18 h) to the distance *d*_*i*_ = |〈PC*i*〉_EMT,0h_ − 〈PC*i*〉_EMT,18h_| between the PC means at 0 h and 18 h (describing EMT-associated change) where *i* = 1 … 22 (Figure S1F). We found that the distance exceeded the standard deviations only for PC1, further indicating that PC1 reflects most of the EMT-associated shape variation in our dataset.

Together, our morphometric analysis yields a high-fidelity and holistic representation of the 3D shapes of cells undergoing EMT. Our analysis further identifies PC1, which encodes multiple cell shape features associated with cell spreading, as a parameter that can be used as a one-dimensional mathematical descriptor of EMT-associated cell shape change.

### Stochastic inference analysis of morphospace trajectories reveals the morphogenetic landscape and cell shape noise dynamics underlying EMT

We then used our morphospace analysis to ask if the shape trajectories of individual cells during EMT could be described as a transition between cell shape attractors in a morphogenetic landscape (Figure 3A). Visual analysis of morphospace cell shape trajectories (Figures 3B, S2I and S2J) suggested that between 3 h and 12 h, coinciding with the time of cell spreading (Figures 2C and 2D), individual cells displayed high variation in their morphospace position, indicative of enhanced cell shape fluctuations (Figures 3 and S3). To further explore these fluctuations, we asked if cell shape dynamics during EMT could be described as a stochastic process driven by deterministic and random forces. We focused on the dynamics of PC1 as a one-dimensional descriptor of EMT-associated cell shape changes, and considered a stochastic process described through the following Langevin equation:

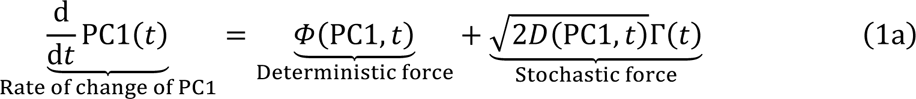

written in the Itô convention^36^, where Γ(*t*) is a Gaussian white noise fulfilling:

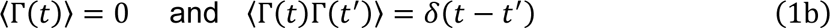

**Figure 3.**
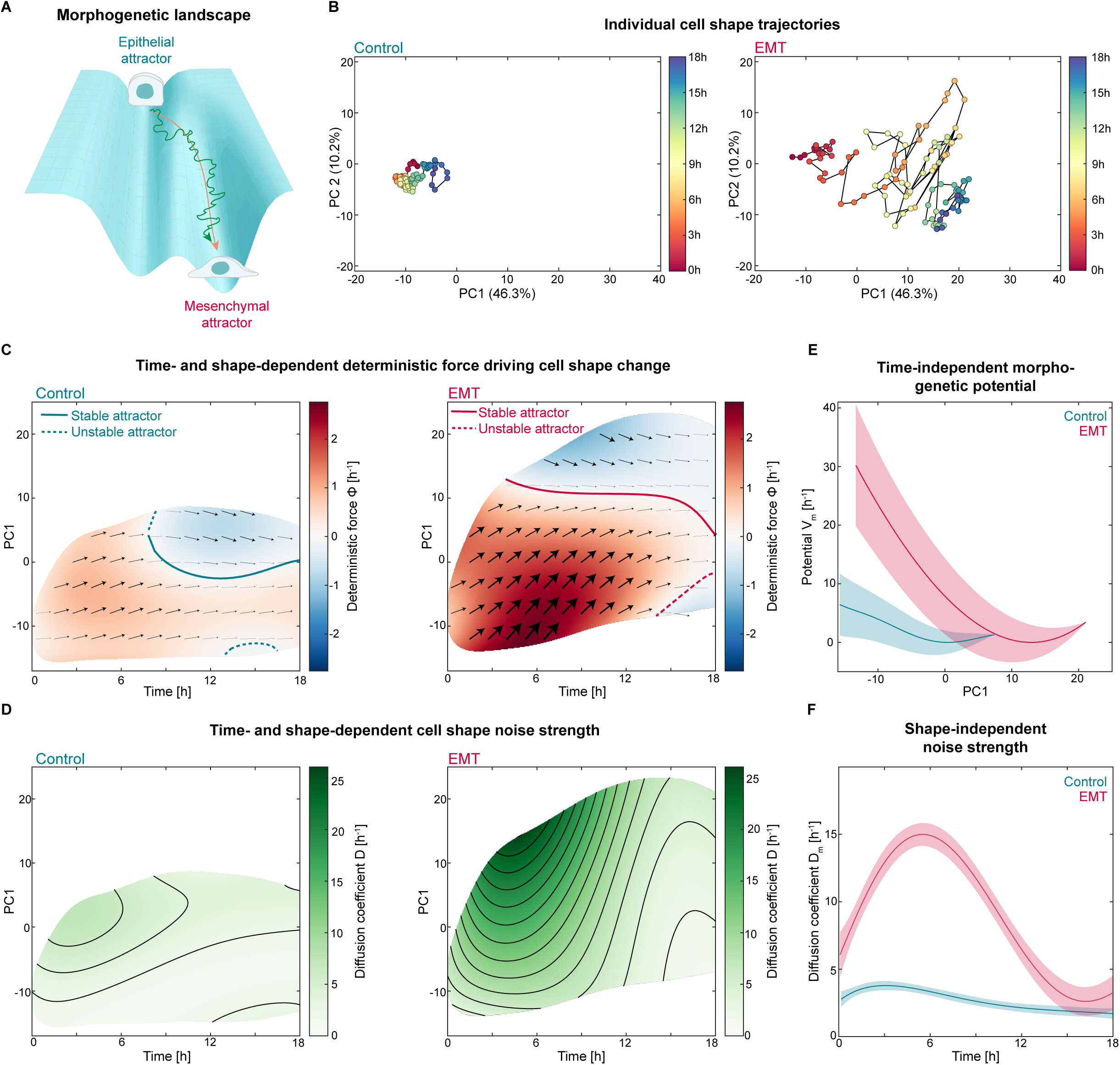
Stochastic inference analysis of morphospace trajectories reveals cell shape attractors and a peak in cell shape noise strength during EMT. (A) Illustration of a morphogenetic landscape. The landscape dictates changes in cell shape, with minima corresponding to local attractors towards which cell shapes converge. The deterministic forces in Equation 1 mediate a persistent trend along the morphogenetic landscape towards attractors (orange trajectory), while contributions due to the stochastic forces in Equation 1 mediate a jittery movement (green trajectory). (B) Individual cell shape trajectory of a control and an EMT cell mapped into the PC1-PC2 morphospace. See Figures S2I and S2J for more trajectories. (C) Deterministic force field 𝛷(PC1, *t*) (Equation 1) inferred from morphospace trajectories of control and EMT cells. The arrows indicate how PC1 changes with time. The curves are illustrations of attractors characterized by Φ = 0, and either 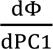 < 0 (for stable attractors) or 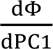 > 0 (for unstable attractors). (D) Diffusion coefficient *D*(PC1, *t*) (Eq. 1) inferred from morphospace trajectories of control and EMT cells. Larger values correspond to larger cell shape noise strength. (E) Morphogenetic potential *V*_*m*_(PC1) of the minimal stochastic model (Equation 2) inferred from morphospace trajectories of control and EMT cells. A minimum in the potential corresponds to a shape attractor. The curves are the best fit and the errorbars indicate the 95% prediction intervals. (F) Diffusion coefficient *D*_*m*_(*t*) of the minimal stochastic model (Equation 2) inferred from morphospace trajectories of control and EMT cells. The curves are the best fit and the errorbars indicate the 95% prediction intervals. (C-F) For control cells N=6, n=102. For EMT cells N=10, n=145.

In these equations, 𝛷 represents the deterministic morphospace force originating from an underlying morphogenetic landscape (Figure 3A); this force drives cell shape towards cell shape attractors. The force 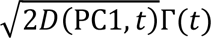 accounts for the stochastic dynamics of cell shape changes, where the diffusion coefficient *D* reflects the strength of cell shape noise in individual cells, which originates from fast processes that are not resolved spatially or temporally but effectively generate forces driving undirected cell shape changes. In this mathematical framework, cell shape fluctuations result from the local structure of the morphogenetic landscape and from the strength of cell shape noise. Of note, the morphospace forces do not have the dimension of mechanical forces (but units of [1/h]), but they reflect the action of mechanical forces that drive cell shape change.

To infer the deterministic and stochastic forces from our experimental data, we adapted a method known as *stochastic (force) inference*^37^ and expanded it to allow for a time dependence of the deterministic and stochastic force fields (see Methods and Supplementary Information). Indeed, in the most generic description, both the deterministic force 𝛷 and the diffusion coefficient *D* can depend on both PC1 and time (Equation 1a). Stochastic force inference analysis of the one-dimensional PC1 trajectories yielded the deterministic force fields 𝛷 (Figure 3C) and diffusion coefficients *D* (Figure 3D) for control and EMT cells. We tested whether the inferred dynamics provide an accurate representation of the experimentally measured PC1 stochastic dynamics by numerically reconstructing PC1 trajectories from the inferred 𝛷(PC1, *t*) and *D*(PC1, *t*). We found that these *in silico* trajectories and their stochastic dynamics exhibited excellent agreement with our experimental observations (Figures S3A-S3F; Methods). This confirmed that our stochastic analysis pipeline reliably extracts the deterministic and stochastic forces underlying experimental cell shape trajectories.

We then analyzed the deterministic force 𝛷 for control and EMT cells (Figure 3C). The magnitude of 𝛷 was generally larger for EMT cells than for control cells, consistent with PC1 remaining mostly unchanged in control cell shape trajectories (Figure S3A). Nonetheless, for control cells after the first 6 hours, we could identify a stable attractor point around PC1 ≈ 0 (solid blue line in Figure 3C), indicative of a shape attractor associated with the epithelial cell state. In contrast, for EMT cells, a stable attractor could be observed as early as 3 h after EMT induction, with the deterministic force pushing cell morphologies towards a stable point at PC1 ≈ 10 (solid red line in Figure 3C). Hence, PC1 ≈ 10 can be interpreted as a shape attractor associated with the mesenchymal state.

We next analyzed the diffusion coefficient *D*, characterizing the strength of the noise acting on individual cell shape trajectories, for control and EMT cells (Figure 3D). We found that EMT cells displayed an overall larger diffusion coefficient *D* than control cells and that for EMT cells, *D* increased with increasing PC1. Furthermore, we observed that *D* exhibited non-monotonous temporal behavior for EMT cells: across PC1 values, *D* first increased with time, reached a maximum around 6 h, corresponding to the mid-timepoint of cell spreading (Figure 2D), and then decreased. This observation is consistent with the high fluctuations in morphospace position observed in the EMT shape trajectories between 3 h and 12 h (Figures 3B and S2J). Finally, we note that multiplicative noise can in principle shift the most likely morphospace position of the shape descriptor (PC1) at steady state away from a deterministic attractor^38^. However, we found that this effect is quantitatively small and thus does not affect our analysis (Figure S4; Methods; Supplemental Information).

Taken together, our stochastic dynamics analysis of shape trajectories shows that EMT is associated with changes in the morphogenetic landscape underlying cell shape, reflecting the existence of an epithelial and a mesenchymal shape attractor. Furthermore, our data indicate that EMT shape change is accompanied by a peak in cell shape noise strength around the time of cell spreading.

### A minimal stochastic model recapitulates cell shape attractors associated with EMT

To simplify the comparison of cell shape trajectories between different conditions, we next considered a minimal stochastic model of EMT, governed by the following equation:

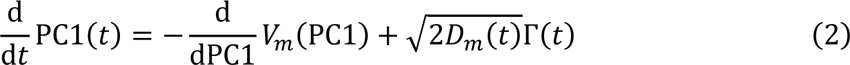

where we introduced the morphogenetic potential *V*_*m*_:

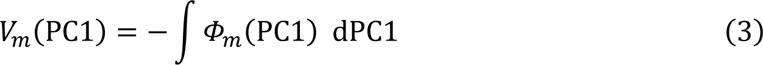

derived from a deterministic force field 𝛷_*m*_(PC1). Equation 2 is equivalent to Equation 1, with the simplifications that the deterministic force is no longer time-dependent but only depends on PC1, and that the diffusion coefficient *D*_*m*_(*t*) only depends on time and not on PC1. We found that the morphogenetic potential *V*_*m*_ inferred with this minimal model displayed a single minimum for both control cells and cells undergoing EMT (Figure 3E). The potential minima, reflecting stable shape attractors, were located at PC1 ≈ 0 and PC1 ≈ 10 for control and EMT cells, respectively, in agreement with the shape attractors inferred with Equation 1 (Figure 3C). The inferred temporal dynamics of the diffusion coefficient *D*_*m*_ further showed that the strength of cell shape noise dramatically increased during EMT and peaked around 6 h (Figure 3F). Finally, we checked that the peak in *D*_*m*_ is not just a feature of PC1 but could also be observed when considering shape trajectories in a high-dimensional space consisting of the first 22 principal components (Figure S3G; Methods). Our analysis shows that even a minimal stochastic model recapitulates the main features of EMT morphodynamics (Figures 3C and 3D), namely the existence of stable epithelial and mesenchymal shape attractors (Figure 3E), and a peak in the strength of cell noise around the time of cell spreading (Figure 3F). We thus used this minimal model to investigate the function of cell shape noise in the rest of the study.

### A theoretical model shows that a transient peak in noise facilitates transitions between attractors

We next asked if the increased shape noise strength we observed during EMT-associated cell spreading could facilitate the cell shape transition. We first approached the question from a theory perspective and investigated the dynamics of a transition between attractors by a particle subjected to a time-dependent noise. We built a mathematical model describing a particle with position *ρ* (analogous to a cell’s morphospace coordinates), transitioning from an epithelial attractor to a mesenchymal attractor. *ρ* starts at an initial position *ρ*_*ep*_, which corresponds to the epithelial shape attractor. The dynamics of *ρ* is governed by a simple quadratic potential 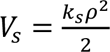 (with the potential strength *k_s_*). The potential minimum corresponds to the mesenchymal attractor, located at *ρ* = 0 (Figure 4A). The dynamics of *ρ* is additionally affected by noise and the time-dependent noise strength reads:

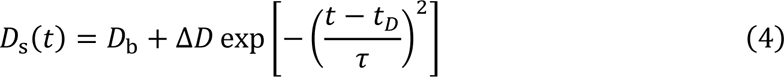

which describes a noise profile with a background noise strength *D*_b_ and a noise peak of height Δ*D* at time *t*_*D*_ and peak width *τ*. We considered three cases (Figure 4B): a noise peak, which qualitatively recapitulates the experimentally observed EMT noise profile (Figure 3F), a constant low noise profile, and a constant high noise profile. The model parameters and mathematical solution are detailed in the Methods and Supplemental Information.

**Figure 4:**
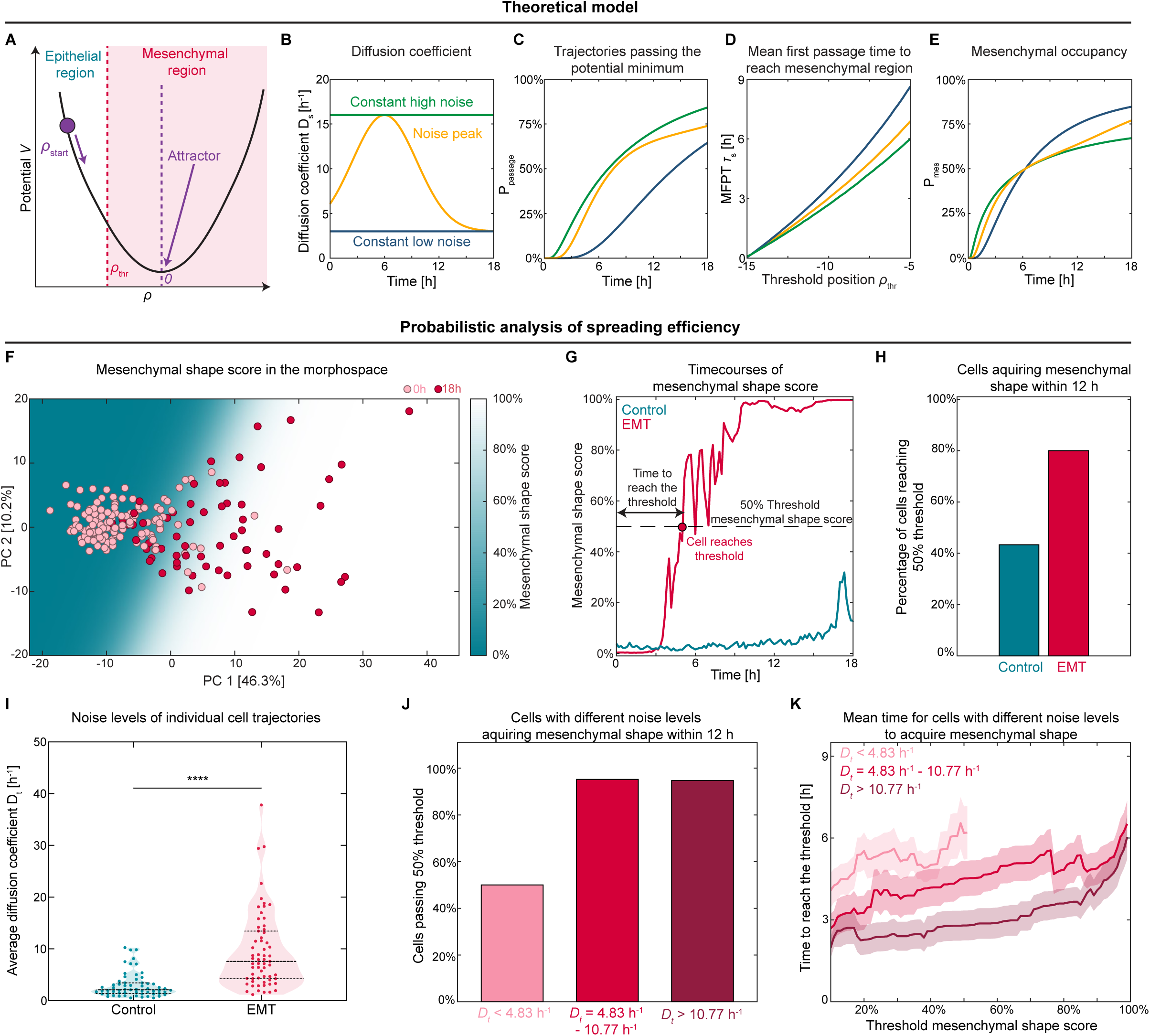
Relationship between cell shape noise strength and the efficiency of EMT-associated cell spreading. (A) Illustration of the stochastic dynamics model for a particle with position *ρ* in a monostable potential *V*_*s*_ = *k*_*s*_*ρ*^2^/2, starting at the initial position *ρ*_ep_ < 0. A threshold *ρ*_thr_ < 0 separates the epithelial and mesenchymal regions. (B) Illustration of the three cases of the diffusion coefficient *D*_*s*_(*t*) considered in the model (see Equation 4 and Methods). (C) Percentage of particles, *P*_passage_, that passed the potential minimum after a time *t*. (D) Mean first passage time *τ*_s_ of particles passing the mesenchymal threshold as a function of the threshold position *ρ*_thr_. (E) Mesenchymal occupancy (percentage of particles, *P*_mes_, located within the mesenchymal region) as a function of time *t*. In (C-E), blue line: “constant low noise” case, green line: “constant high noise” case, yellow line: “noise peak” case. (F) Probability map of a shape being mesenchymal in the PC1-PC2 morphospace. Color gradient: mesenchymal shape score. Pink and red datapoints: 0 h and 18 h EMT cells, respectively (same dataset as in Figure 2A, S2A and S2B). (G) Mesenchymal shape score of individual representative control and EMT cells as a function of time. (H) Percentage of control (N=8, n=55) and EMT (N=8, n=49) cells that pass a 50% mesenchymal shape score threshold within 12 h. (I) Distribution of average diffusion coefficients for individual cells, *D*_*t*_, for control (N=8, n=61) and EMT (N=8, n=63) cells, (P<0.0001, Mann-Whitney test), dashed line: median, dotted lines: quartiles. (J) Percentage of EMT cells with weak cell shape noise strength (*D*_*t*_ < 4.83 h^−1^), moderate cell shape noise strength (4.83 h^−1^ ≤ *D*_*t*_ ≤ 10.77 h^−1^), and large cell shape noise strength ( *D*_*t*_ > 10.77 h^−1^) that reaches a 50% mesenchymal shape score threshold within 12 h. n=21 for each category. (K) Mean time taken by EMT cells to reach a given mesenchymal shape score threshold; mesenchymal score thresholds higher than 10% (x-axis) are considered to assess a range of spreading extent. Mean ± SEM.

We first compared how many trajectories reached the potential minimum after a given time *t* for the different noise profiles. We found that trajectories with higher noise (noise peak or constant high noise) reached the potential minimum earlier than trajectories with low noise (Figure 4C). In our experiments, mesenchymal cell shapes display a relatively broad distribution around the mesenchymal shape attractor (Figure 2A), reaching the potential minimum itself may thus be an insufficient readout of successful shape transition. We thus analyzed transition dynamics towards a broader mesenchymal region defined by *ρ* > *ρ*_*thr*_, where positions above a threshold *ρ*_*thr*_ are considered mesenchymal (Figure 4A). Trajectory simulations showed that across all thresholds, trajectories with higher noise (peak and constant high noise) reached the mesenchymal regions earlier than trajectories with low constant noise (Figure 4D). Thus, our model indicates that higher noise accelerates the transition towards a new potential minimum.

However, high noise could also drive trajectories away from the new minimum once it has been reached, reducing the robustness of the transition. To test this, we calculated the percentage of trajectories located within the mesenchymal region, or mesenchymal occupancy *P*_mes_, as a function of time (Figure 4E). Initially more trajectories with higher noise reached the mesenchymal region, consistent with high noise accelerating the transition. However, at later timepoints, mesenchymal occupancy became highest for particles with constant low noise, while mesenchymal occupancy for particles with constant high noise stalled. Interestingly, mesenchymal occupancy for particles with a noise peak steadily increased, converging towards the occupancy for constant low noise particles over time. Taken together, our mathematical model suggests that a transient noise peak combines the contrasting benefits of constant low and high noise: the initial higher noise accelerates the transition towards a new attractor (Figure 4C and 4D), while the later decrease in noise makes the transition more robust (Figure 4E).

### Trajectories with higher cell shape noise strength display faster cell spreading during EMT

We then asked how cell shape noise strength affected cell shape transition dynamics in our experiments. As the time of shape transition can be difficult to pinpoint in the noisy cell shape trajectories (Figure 3B), we developed a mathematical framework to define the efficiency of spreading probabilistically. We assumed that cells undergoing EMT displayed epithelial shapes at 0 h and mesenchymal shapes at 18 h post HGF addition, and used our shape data to train a support vector machine classifier for binary classification of epithelial and mesenchymal shapes. We trained the classifier using the (PC1, PC2) coordinates of 0 h and 18 h EMT cells; PC2 was included to increase the accuracy of the predictor, as we observed a weak difference in the ensemble dynamics of PC2 between EMT and control cells (Figure S2G). We then used this classifier to derive the probability map of mesenchymal shapes, which assigns a mesenchymal shape score to every location in the PC1-PC2 morphospace (Figure 4F) and tracked how this score changed for individual cells undergoing EMT (Figure 4G). Finally, we used the resulting mesenchymal shape score timecourse to quantify, for each threshold mesenchymal shape score, a shape transition time as the time taken by a cell to reach the threshold (Figure 4G), and the percentage of cells that reach the threshold within 12 h (Figure 4H). The interval of 12 h was selected as a cutoff, as it corresponds to the time window of cell spreading (Figure 2C). As expected, we found that a larger fraction of EMT cells reached any given threshold mesenchymal shape score compared to control cells (Figures 4H and S5A). Together, our data-driven method allows for quantification of the efficiency of EMT-associated cell spreading through the timing and extent of the cell shape change.

We then asked how cell spreading efficiency related to cell shape noise strength for individual cells. To characterize noise strength in single cells, we computed an average diffusion coefficient *D*_*t*_for each cell morphospace trajectory (Methods). Consistent with our stochastic inference analysis (Figure 3F), we found that control cells had lower *D*_*t*_ than EMT cells (Figure 4I). We next grouped EMT cell shape trajectories into three equally sized populations, displaying low, moderate, or large cell shape noise strength. Interestingly, we found that cells with strong and moderate noise strength were more likely to cross any mesenchymal shape score threshold than cells with weak noise strength (Figure 4J and S5B). Moreover, the larger the shape noise, the earlier cells reached mesenchymal shapes across all mesenchymal shape score thresholds (Figure 4K). These observations are in agreement with our theoretical model predictions that higher noise increases transition efficiency (Figure 4C and 4D). Together, our data show that higher cell shape noise correlates with faster and more extensive cell spreading during EMT.

### Experimentally reducing cell shape noise delays EMT-associated cell spreading

Finally, we asked how reducing cell shape noise experimentally affects cell shape transition dynamics. We tested if perturbing the mechanical balance at the cell surface might affect cell shape noise during EMT. Cell spreading requires an increase in actin protrusivity and a reduction in plasma membrane tension to accommodate increased protrusion formation^39–41^. We thus sought to perturb actin and membrane dynamics, and assessed effects on cell shape noise strength and spreading efficiency. We first interfered with actin protrusivity by inhibiting the activity of the small GTP-ase Rac1, a key regulator of protrusive actin polymerization in EMT^32,42^. Upon Rac1 inhibition with the small molecule inhibitor NSC23766^43^, cells displayed considerably reduced spreading 18 h after EMT induction compared to untreated EMT cells (Figure 5A, Movie S4). Morphospace analysis further demonstrated that cells with reduced Rac1 activity generally displayed less spread cell shapes characterized by lower PC1 (Figures 5B, 5C and S5C). Stochastic inference showed that Rac1 inhibition in EMT cells shifted the shape attractor to lower PC1 values, and dramatically decreased cell shape noise strength (Figure 5D). We then analyzed cell shape spreading efficiency from the mesenchymal shape score timecourses (Figure S5D). We found that Rac1 inhibition drastically reduced the percentage of cells that reached any mesenchymal shape threshold after 12 h (Figures 5E and S5E), consistent with the overall reduction in cell spreading (Figures 5B and 5C). Interestingly, the cells that did display spreading reached their spread shapes significantly later than untreated ЕМТ cells across all mesenchymal shape scores (Figure 5F).

**Figure 5:**
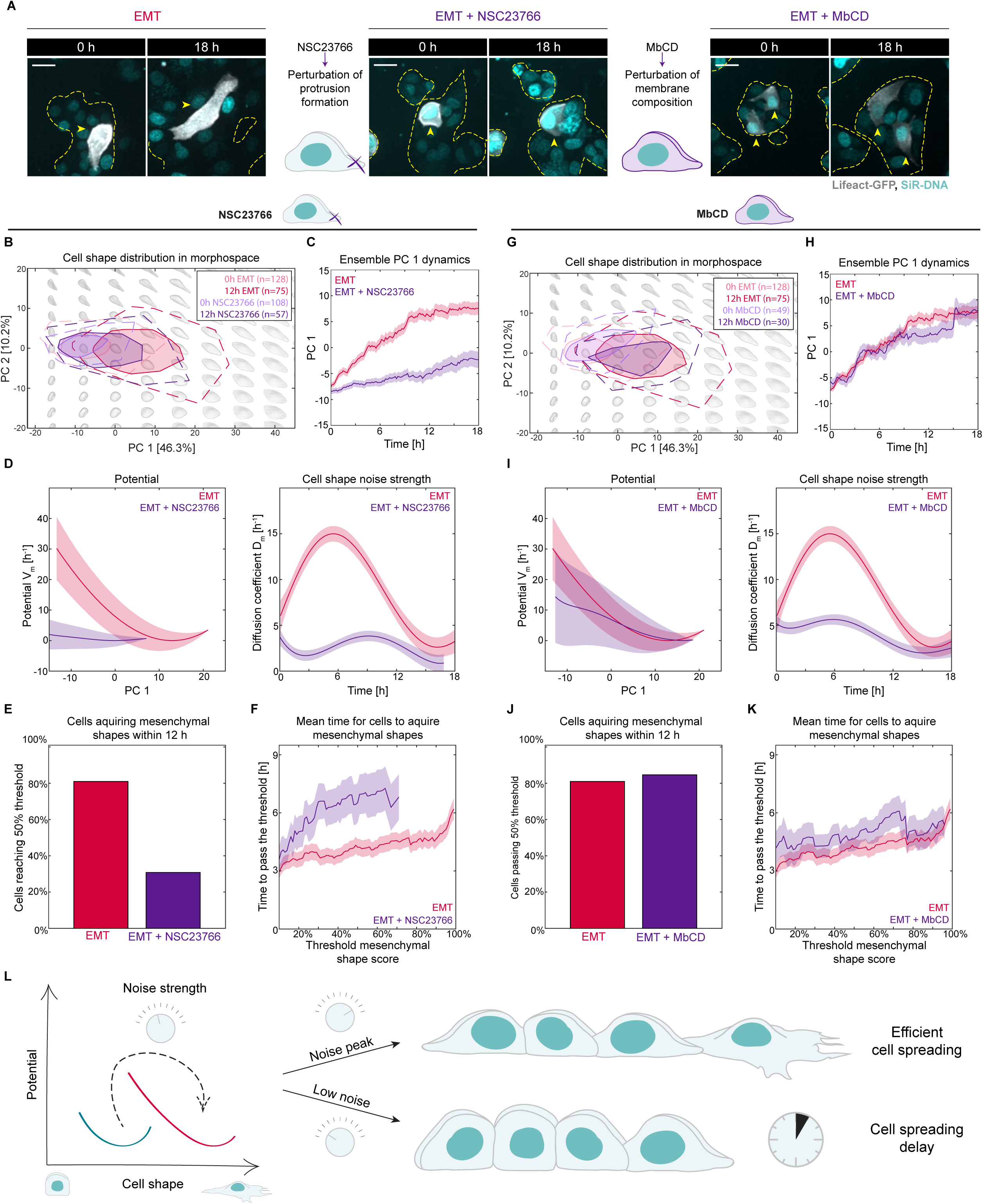
Reducing cell shape noise strength interferes with EMT-associated cell spreading. (A) Representative images at 0 h and 18 h after EMT induction of unperturbed EMT cells (left) and cells treated with 100 µM Rac1 inhibitor NSC23766 (middle) or 1 mM MbCD (right). Labels: Lifeact-GFP (F-actin, gray) and SiR-DNA (nuclei, cyan). Yellow dashed lines: colony outlines, arrowheads: representative Lifeact-GFP cells. Scale bars: 20 µm. (B) Bagplot representation of the distribution of untreated (N=10, n=128 and N=10, n=75) and NSC23766-treated EMT cells (N=7, n=108 and N=6, n=57) in the morphospace at 0 h and 12 h. Background gray shapes: reconstructed theoretical cell shapes for different combinations of PC1 and PC2. (C) Ensemble dynamics of PC1 of untreated (N=10, n=145) and NSC23766-treated (N=7, n=115) EMT cells. Mean ± SEM. (D) Morphogenetic potential *V*_*m*_(PC1) and diffusion coefficient *D*_*m*_(*t*) of untreated and NSC23766-treated EMT cells inferred from their PC1 morphospace trajectories. *V*_*m*_(PC1) and *D*_*m*_(*t*) curves are the best fit and error bars indicate the 95% prediction intervals. (E) Percentage of untreated (N=10, n=63) or NSC23766-treated (N=6, n=52) EMT cells that pass a 50% mesenchymal shape score threshold within 12 h as a function of the. (F) Mean time taken by untreated or NSC23766-treated EMT cells to reach a given mesenchymal shape score threshold. Mean ± SEM. (G) Bagplot representation of the distribution of untreated (N=10, n=128 and N=10, n=75) and MbCD-treated EMT (N=4, n=49 and N=4, n=30) cells in the morphospace at 0 h and 12 h. Background gray shapes: reconstructed theoretical cell shapes for different combinations of PC1 and PC2. (H) Ensemble dynamics of PC1 of untreated (N=10, n=145) and MbCD-treated (N=4, n=54) EMT cells. Mean ± SEM. (I) Morphogenetic potential *V*_*m*_(PC1) and diffusion coefficient *D*_*m*_(*t*) of untreated and MbCD-treated EMT cells inferred from their PC1 morphospace trajectories. *V*_*m*_(PC1) and *D*_*m*_(*t*) curves are the best fit and error bars indicate the 95% (J) Percentage of untreated (N=10, n=63) or MbCD-treated (N=4, n=26) EMT cells that pass a 50% mesenchymal shape score threshold within 12 h. (K) Mean time taken by untreated or MbCD-treated EMT cells to reach a given mesenchymal shape score threshold. Mean ± SEM. (L) Schematic summary of the effect of cell shape noise on transition dynamics between cell shape attractors. Transient increase in cell shape noise leads to efficient cell spreading while low cell shape noise levels delay the cell shape transition.

We then treated cells with methyl-β-cyclodextrin (MbCD), a cholesterol-depleting compound that has been shown to increase membrane tension in various cell types^44,45,41^, including MDCK cells^46^. While some MbCD-treated cells displayed reduced spreading in EMT, many cells displayed spreading visually comparable to untreated cells 18 h after EMT induction (Figure 5A, Movie S4). Morphospace analysis demonstrated that MbCD-treated cells displayed similar values of PC1 as untreated EMT cells (Figure 5G). The ensemble EMT spreading dynamics were also similar in MbCD-treated and untreated EMT cells (Figures 5H and S5C). Stochastic inference further showed that the morphogenetic potential of MbCD-treated EMT cells was comparable to that of untreated EMT cells, with a minimum around PC1 = 10 (Figure 5I). Strikingly, however, MbCD treatment led to a strong reduction in cell shape noise strength (Figure 5I). We then analyzed individual cell spreading efficiency from the mesenchymal shape score timecourses (Figure S5D). We found that the percentage of cells acquiring mesenchymal shapes after 12 h was only weakly reduced compared to untreated cells (Figures 5J and S5E), in agreement with the comparable morphogenetic potentials of MbCD-treated and untreated EMT cells (Figure 5I). However, MbCD treatment led to a ∼20% increase in the time required to reach a mesenchymal threshold compared to untreated EMT cells across all mesenchymal cell shape scores (Figure 5K and S5E). These findings agree with our theoretical prediction that decreasing cell shape noise strength could slow down the transition from one shape attractor to another (Figure 4D). Together, our theoretical model and perturbation experiments strongly suggest that the peak in cell shape noise observed during EMT plays a functional role, accelerating EMT-associated cell spreading (Figure 5L).

## DISCUSSION

In this study, we investigated cell shape dynamics during EMT-associated cell spreading. We employed a data-driven approach to quantify the morphodynamics of 3D cell shape changes of MDCK cells undergoing EMT. Most studies of morphogenesis use intuitive shape measures (e.g., volume, solidity, aspect ratio) as quantitative features describing cell morphology^11,16,47^. However, such features are often subjectively chosen and can thus be biased. Furthermore, they do not constitute unique descriptors, and thus cannot be used to directly reconstruct the original cell shapes. To avoid these limitations, we used spherical harmonics decomposition, a mathematical framework that has been recently attracting increasing attention for cell shape representation^19,22–26^. For unbiased and interpretable dimensional reduction, we employed PCA, as it represents low-dimensional features as linear combinations of the high-dimensional spherical harmonics descriptors. The linearity of the PCA mapping allows for defining distances between distinct shapes, and for the reconstruction of theoretical shapes corresponding to each position of the PCA morphospace (see grey shapes in e.g. Figure 2A). Together, this analysis enabled the representation of complex 3D shapes and shape dynamics of individual cells in a low-dimensional morphospace in an objective and interpretable way.

We then used mathematical modeling to analyze the stochastic dynamics of EMT cell shape trajectories in morphospace. Our stochastic model shows that EMT-associated shape changes can be described using the mathematical framework of attractors (Figure 3). How EMT-associated signaling pathways regulate the epithelial and mesenchymal shape attractors we identify here will be an interesting avenue for future studies. It will also be interesting to investigate to what extent mechanical constraints on the organization of cellular components controlling cell surface mechanics contribute to determining cell shape attractors. Recent work suggests that actin networks can only display a limited number of configurations, which could reflect constraints on the wiring of upstream signaling pathways^13^, but could also reflect mechanical and structural constraints on the configurations an actin network can display^48,49^. It is tempting to speculate that such molecular and mechanical constraints could drive the emergence of attractors in actin networks organization, which could in turn determine attractors in cell shape.

Strikingly, we found that the transition between shape attractors is associated with a peak in cell shape noise strength. Importantly, our stochastic inference analysis reveals a peak in noise at the individual cell level, not a collective effect reflecting, e.g., asynchrony or heterogeneity of shape transition dynamics between cells. Our model and perturbation experiments strongly suggest that noise functionally facilitates cell shape transition during EMT (Figure 5L). Noise has been shown to be a driver of transitions between attractors in quantum optics, chemical reaction kinetics and population dynamics^50,51^. In genomics, noise stemming from fluctuations in the expression and interactions of genes and proteins^52–54^ has been shown to potentially affect transitions, as noise can distort the underlying epigenetic landscape, making it an essential regulator of cell state^55^. Though cell shape noise has been less explored, it has been proposed to play a functional role in driving chemotactic response in neuronal growth cones^56^ and cell sorting during embryonic development^57^. Building on our findings, it will be exciting to investigate to what extent cell shape noise also contributes to other types of cellular shape transitions, and how shape noise is regulated.

The coupling between cell shape, the underlying mechanics, and cell state has recently been attracting increasing attention^41,58,17^. In several systems, cell shape change has been shown to be a potent correlate and predictor of cell fate^59,26^. The morphospace analysis we developed here provides a highly sensitive method for cell shape quantification, and could thus constitute a powerful tool for non-destructive monitoring of cellular transitions in other contexts. Furthermore, our mathematical analysis framework could be expanded to include other cellular features, such as the levels and distributions of state markers. Such multi-dimensional analysis will constitute a powerful investigation tool that could help establishing a more holistic definition of what constitutes a cell state^60^.

Finally, we found that modulating the strength of cell shape noise can influence the rate of EMT-associated cell spreading without altering the morphogenetic landscape and, thus, contribute to regulating the timing of the process. Mechanisms to adjust the timing of cell spreading could be functionally essential during EMT in physiological processes like wound healing and development. Delayed wound closure increases the risk of infections^61^. In development, cell shape and fate changes need to be coordinated in time and space. For instance, in zebrafish gastrulation the timing of gastrulating cell specification is essential for coordinating cell protrusivity and internalization capacity while preserving tissue patterning^62^. In both contexts the exact timing of cell shape change may be key for organism survival. Our work suggests the intriguing possibility that cells could tune the levels of cell shape noise to flexibly adjust the timing of cellular shape transitions.

## RESOURCE AVAILABILITY

This study did not generate new unique reagents. All data and MATLAB scripts are available upon request.

## ACKNOWLEDGEMENTS

We would like to thank Margherita Battistara, Kevin Chalut, Joe Howard, Laura Machesky Xiaoyi Ouyang, Ana Raffaelli, Aki Stubb, Marta Urbanska for feedback on the manuscript, Nir Gov and Rutger Hermsen for helpful discussions about the data analysis and theoretical modelling. We thank Lakshmi Balasubramaniam for help with hydrogel protocol optimization. We also thank Windie Höfs and Guillaume Charras for providing the MDCK cell lines and input on cell culture conditions. We thank the Cambridge Advanced Imaging Centre for assistance with microscopy.

We also thank the Paluch lab (in particular Margherita Battistara, Will Foster, Tasmin Sarkany, Belle Sow, Aki Stubb, Baptiste Vauléon) and the Salbreux lab (in particular Silvia Grigolon, Diana Khoromskaia, Quentin Vagne, Max Kerr Winter) for discussions and feedback throughout the project. We thank Fiona Morgan, Liz Williams and Alex Winkel from the Paluch lab for technical support.

E.K.P. acknowledges funding from the European Research Council (Consolidator Grant 820188-NanoMechShape), Medical Research Council UK (MRC Programme Award MC_UU_12018/5), and the Philip Leverhulme Trust (Leverhulme Prize in Biological Sciences to E.K.P.). W.P. acknowledges financial support from the Herchel Smith Fund (Herchel Smith Postdoctoral Fellowship).

## AUTHOR CONTRIBUTIONS

E.K.P., G.S., I.Y. and W.P. developed the project idea; I.Y. developed and performed the wet lab experiments; I.Y. and W.P. conducted the spinning disk microscope imaging; I.Y. and W.P. conducted the segmentation; W.P. was responsible for the data analysis and theoretical modelling; E.K.P., G.S., I.Y. and W.P. prepared the manuscript; E.K.P. and G.S. supervised the project, discussed the data and results.

## SUPPLEMENTARY FIGURE TITLES AND LEGENDS

**Figure S1:**
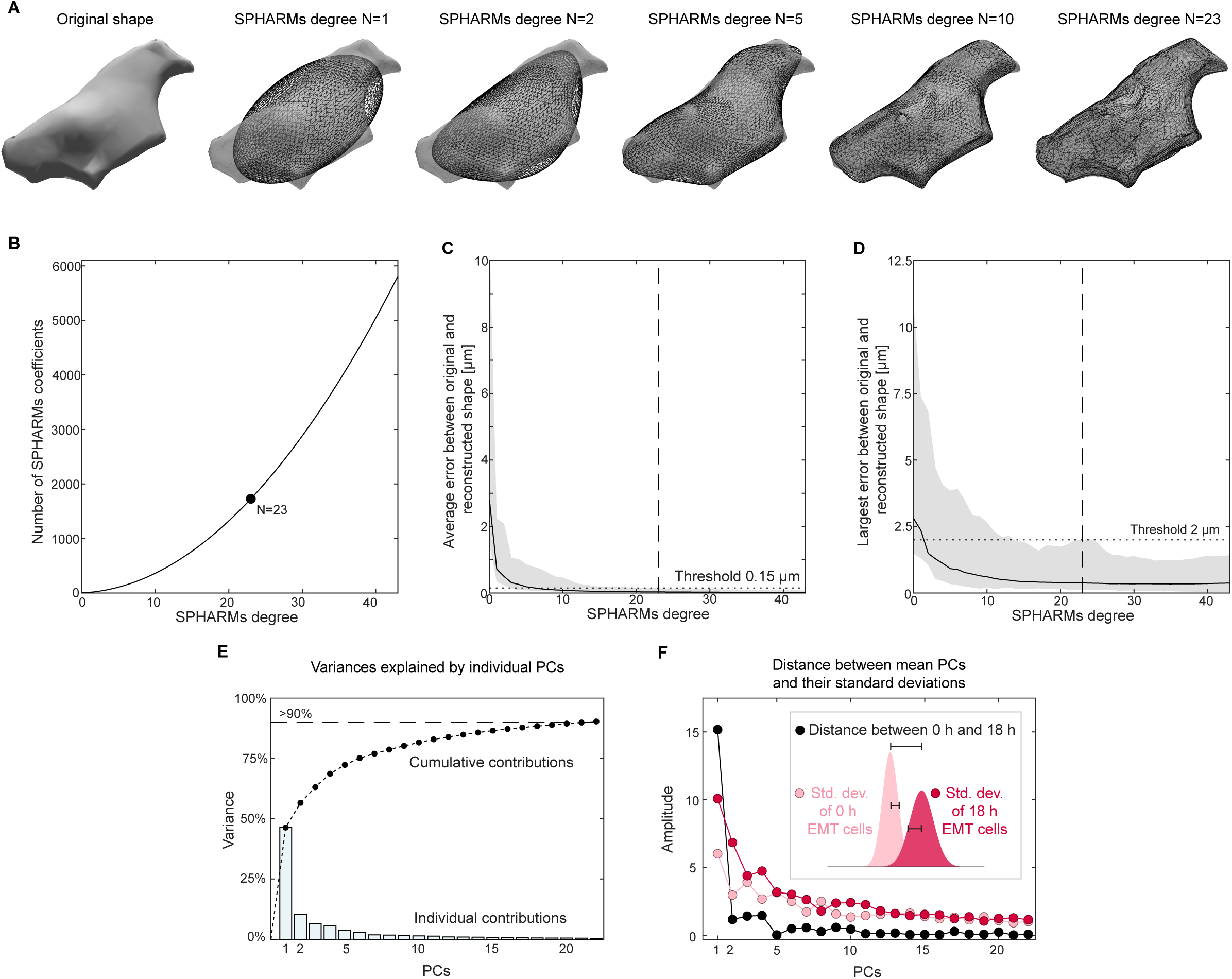
SPHARM reconstruction and identification of most suitable SPHARMs degree, related to Figure 1. (A) Reconstruction of cell shapes from the SPHARMs coefficients of different degrees. The grey cell shows the original cell shape, and the black meshes represent the reconstructed shapes. (B) Number of SPHARMs coefficients as a function of SPHARMs degree *N*, given by 3(*N* + 1)^2^. In this study, we used *N* = 23, corresponding to 1728 coefficients. (C, D) Mean and largest error between original and reconstructed shapes for 0 h and 18 h control and EMT cells (N=10, n=346) as a function of SPHARMs degree. We defined a threshold of 2 µm for the largest error and 0.15 µm for the mean error (dotted lines) as largest acceptable errors to identify the lowest possible SPHARMs degree. We found that this was fulfilled for *N* = 23 (black dashed lines). The error bars indicate the range of error values (lowest and highest maximum and average error of a cell). (E) Individual and cumulative percentage of the total cell shape variance explained by each principal component. (F) Comparison of the distance *d*_*i*_between the average PC*i* at 0 h (N=10, n=128) and at 18 h (N=10, n=63) in EMT cells and of their standard deviations *σ*_*i*,EMT,0h_ and *σ*_*i*,EMT,18h_ for *i* =

**Figure S2:**
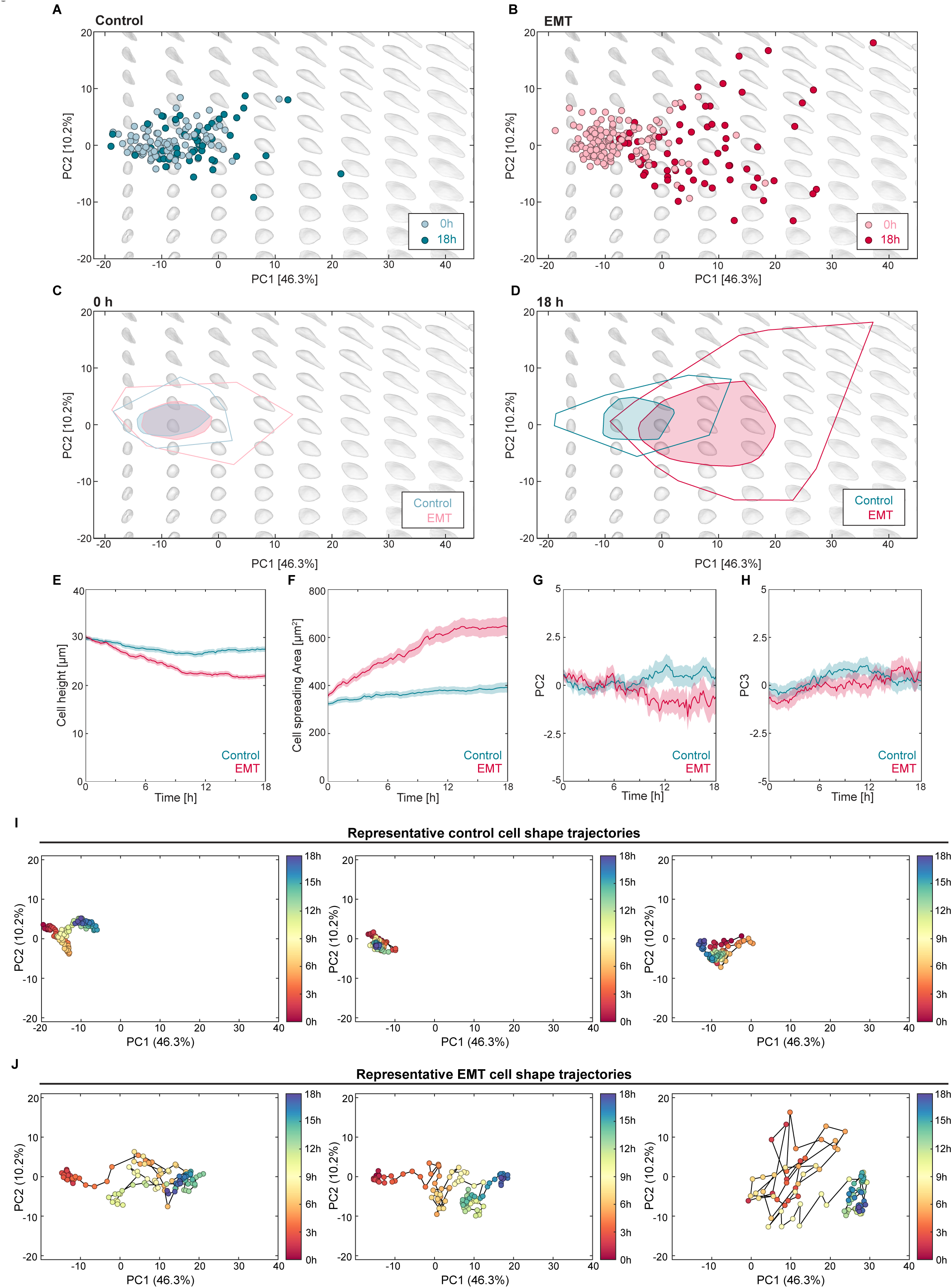
Distribution of control and EMT cells in the morphospace and cell shape dynamics during EMT, related to Figure 2 and 3. (A) Point cloud of all control cells in morphospace at 0 h (N=6, n=100) and 18 h (N=6, n=55). (B) Point cloud of all EMT cells in the morphospace 0 h (N=10, n=128) and 18 h (N=10, n=63) after EMT induction. (C) Bagplots of control and EMT cells in morphospace 0 h after EMT induction. Bagplots are a higher-dimensional expansion of 1d boxplots and used here to visualize the range of a point cloud distributions in the PC1-PC2 morphospace. See Methods for details on the bagplots. (D) Bagplots of control and EMT cells in morphospace 18 h after EMT induction. (A-D): Gray background shapes: reconstructed shapes corresponding to the specific location in the PC1-PC2 morphospace. (E,F) Ensemble dynamics of cell height (E) and cell projected area (F) of EMT cells (N=12, n=145) and control cells (N=8, n=102). Mean ± SEM. (G,H) Ensemble dynamics of PC2 (G) and PC3 (H) of EMT cells (N=12, n=145) and control cells (N=6, n=102). Mean ± SEM. (I) Representative cell shape trajectories of control cells mapped into the morphospace. (J) Representative cell shape trajectories of an EMT cells mapped into the morphospace.

**Figure S3:**
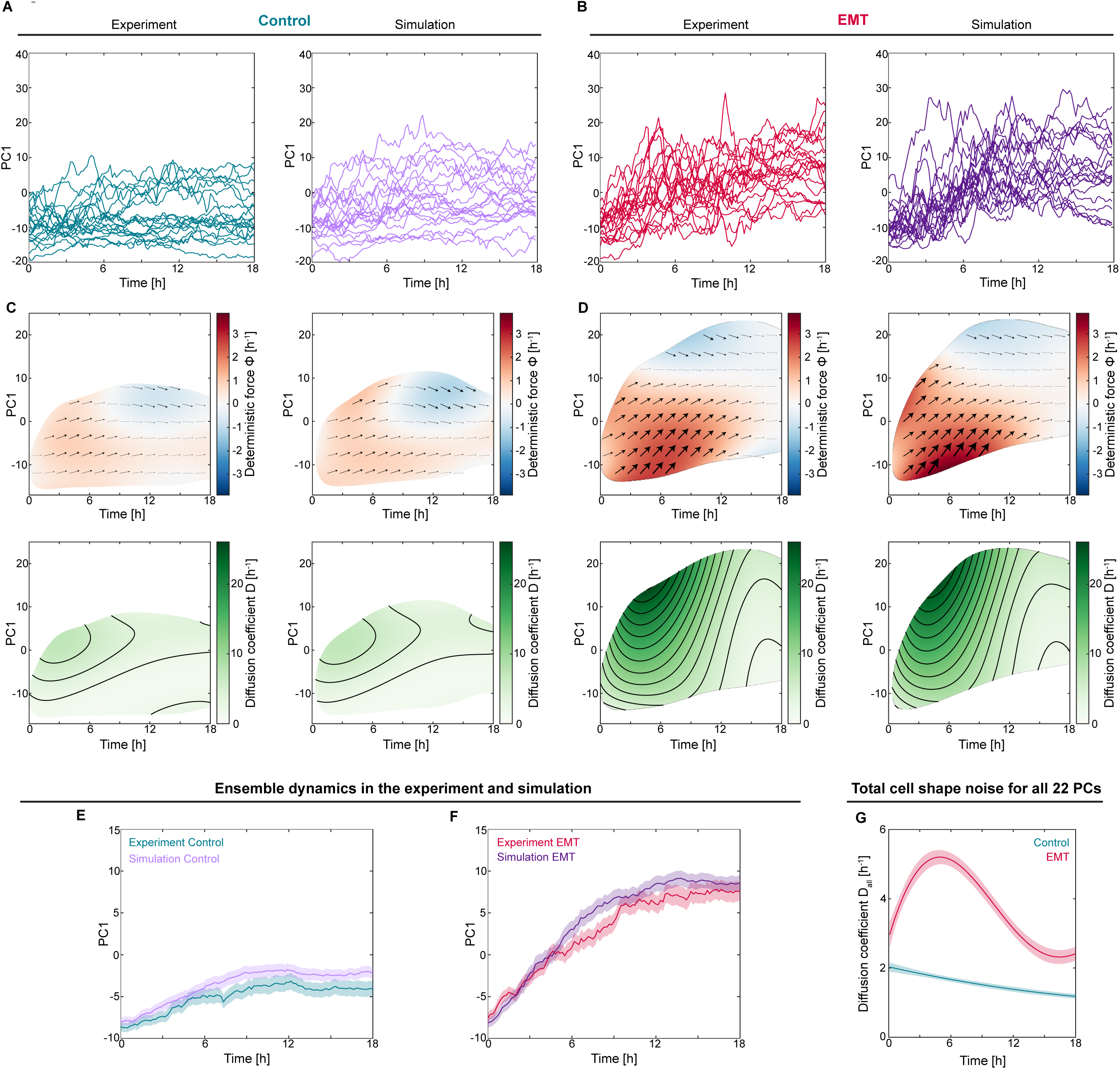
Verification of stochastic inference approach, related to Figure 3. (A,B) PC1 trajectories of experimental control (A) and EMT (B) cells and *in silico* trajectories for the deterministic force field and diffusion coefficient inferred through our stochastic dynamics analysis (Figures 3C and 3D; Equation 1). Details of the generation of *in silico* trajectories is provided in Methods. (C,D) Deterministic force field 𝛷(*ρ*, *t*) and diffusion coefficient *D*(*ρ*, *t*) of Equation 1 inferred from experimental and simulated PC1 data for control (C) and EMT (D) cells. The plots show good agreement between experiments and simulations. (E,F) Ensemble dynamics of experimental (N=6, n=102 and N=12, n=145) and simulated PC1 trajectories (n=100) of control and EMT cells exhibit good qualitative agreement. Mean ± SEM. (G) Total diffusion coefficient extracted through stochastic inferrence analysis of the first 22 principal components combined (Methods).

**Figure S4:**
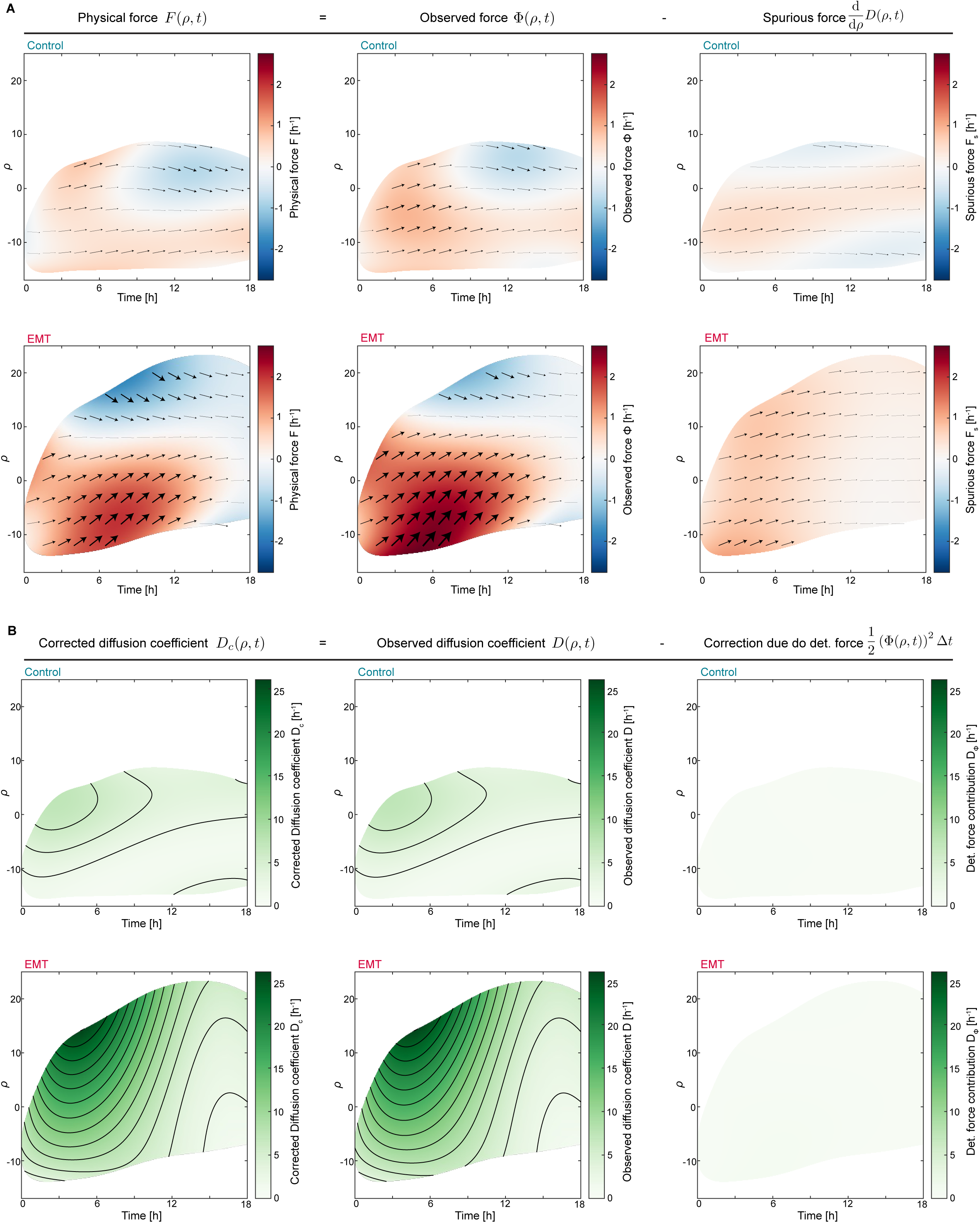
Contributions due to multiplicative noise to 𝚽 and deterministic displacements to 𝑫 have no qualitative impact on the stochastic inference analysis of EMT cell shape trajectories, related to Figure 3. (A) Contribution due to a spurious force d*D*(*ρ*, *t*)/d*ρ* (Supplemental Information) to the inferred deterministic force Φ(*ρ*, *t*) for control and EMT cells. We found that subtracting the spurious force from the inferred deterministic force field did not lead to any qualitative differences between Φ(*ρ*, *t*) and the corrected physical force F(*ρ*, *t*). (B) Contributions of the deterministic force field Φ(*ρ*, *t*) to the diffusion coefficient *D*(*ρ*, *t*) for control and EMT cells. The mean squared displacements mediated by the deterministic force field, given by 1/2(Φ(*ρ*, *t*))^2^ Δ*t* (for a time step Δ*t* = 10 min; Methods; Equation 14) are so small that they can be neglected when calculating the diffusion coefficient *D*.

**Figure S5:**
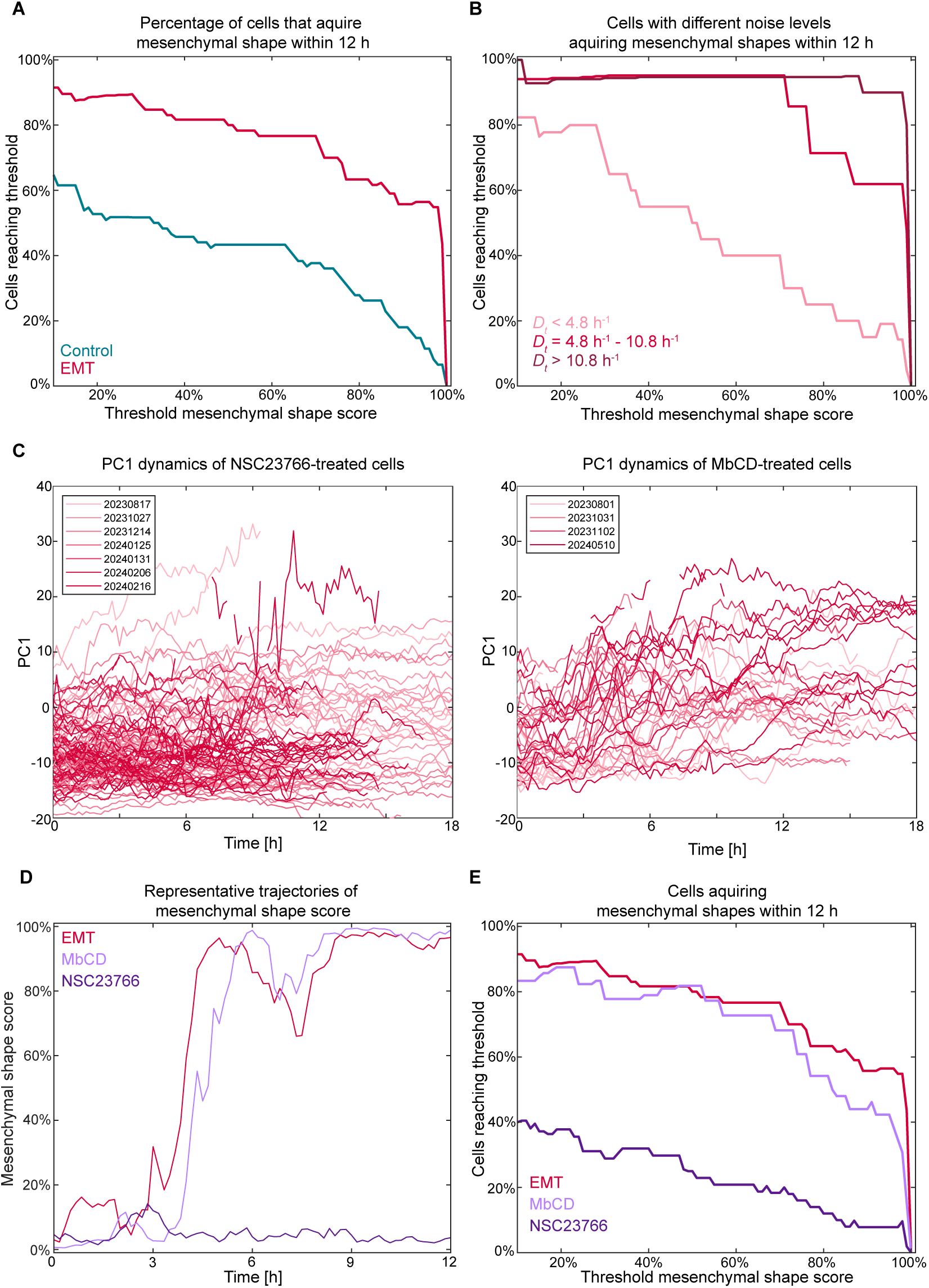
Effect of drug treatments on EMT-associated cell spreading, related to Figure 5. (A) Proportion of control and EMT cells that acquire mesenchymal shape within 12 h post EMT induction for different mesenchymal shape scores. (B) Proportion of EMT cells with different noise levels that acquire mesenchymal shapes within 12 h post EMT induction. (C) PC1 dynamics of individual cells treated with 100 µM Rac1 inhibitor NSC23766 (left) and 1mM MbCD (right). Insets highlight independent experiments. (D) Mesenchymal shape score as a function of time for individual representative untreated cell (red line) and cells treated with 100 µM Rac1 inhibitor NSC23766 (dark purple line) and 1mM MbCD (light purple line). (E) Proportion of untreated EMT cells and EMT cells treated with 100 µM Rac1 inhibitor NSC23766 (dark purple line) and 1mM MbCD (light purple line) that acquire mesenchymal shape within 12 h post EMT induction for different mesenchymal shape scores.

## METHODS

### EXPERIMENTAL MODEL AND STUDY PARTICIPANT DETAILS

Labeled cell lines used in the study are all derived from MDCK-II cells. MDCK-II cells are a spontaneously immortalized cell line originating from adult female Canis lupus familiaris (Dog, breed: Cocker Spaniel) (NCBI Taxonomy: 9615). The cell line has not been authenticated.

### Cell culture

MDCK-II cell lines (wild type (WT) and stable lines expressing Lifeact-GFP or H2B-iRFP) were a gift from Professor Guillaume Charras (London Centre for Nanotechnology, University College London, UK). MDCK cells were cultured at 37°C in a humidified atmosphere of 5% CO_2_. Cells were passaged every 3-4 days using standard cell culture protocols and disposed of after no more than 30 passages. The culture medium was composed of high-glucose Dulbecco’s Modified Eagle Medium, GlutaMAX (Thermo Fisher Scientific, #31966047) supplemented with 10% FBS (Thermo Fisher Scientific, #10500-064), 1% penicillin-streptomycin (Thermo Fisher Scientific, #P4333) and 25 mM HEPES (Thermo Fisher Scientific, #15630056). Cells were tested for mycoplasma monthly and free of contamination.

## METHOD DETAILS

### Hydrogel fabrication

Polycrylamide hydrogels were fabricated as in Ref.^34^ with minor changes. Briefly, support coverslips and cell culture chambers were silanized for two hours with 2% 3-(Trimethoxysilyl)propyl methacrylate (Merck, #M6514) and 1% acetic acid diluted in ethanol, and air-dried in an oven at 60°C for one hour. Hydrogel solutions were prepared to produce gels of 3.5 kPa stiffness (48 µl 40% Acrylamide (Merck, #A4058), 36 µl 2% Bisacrylamide (Merck, #M1533); 20 µl 2 M 6-(prop-2-enoylamino)hexanoic acid (AK Scientific, #6845AL), 389.5 µl MiliQ water, 1.5 µl TEMED (Merck, #T22500), 5 µl 10% APS (Merck, #A3678)) and polymerized between a support coverglass (cell culture chamber or a silanized coverslip) and a clean top coverslip for 25-35 minutes. To ensure that cell morphology was not affected by the underlying glass substrate, gel mix quantity was adjusted depending on coverslip diameter to produce hydrogels with thicknesses above 100 µm. After polymerization, hydrogels were soaked in PBS overnight. The top coverslips were removed on the following day. Hydrogels were then activated by 30 min treatment in 0.2 M EDAC (Merck, #03450) and 0.5 M NHS (Acros Organics, #157272500) in MES buffer, pH 6.1, and coated with 25 µg/ml laminin (Sigma-Aldrich, #L2020) diluted in HEPES buffer (50 mM pH 8.5) overnight at 4°C. After washes, hydrogels were blocked with 0.5 M Ethanolamine (Merck, #411000) in HEPES buffer. Hydrogels were stored in basal medium at 4°C for no more than two weeks.

### Microscope image acquisition

#### Cell plating and preparation

Cells were seeded at sub-confluent levels on 3.5 kPa hydrogels coated with 25 µg/ml mouse laminin (Merck, #L2020) in 2-well chambered coverglasses (Nunc™ Lab-Tek™ II, #155379 or Ibidi µ-Slide, #80287), allowing control and EMT cells to be compared directly. Unlabeled MDCK cells were mixed with Lifeact-GFP MDCK cells (15:1 ratio) enabling the imaging of cell morphology of single labeled cells located in clusters of unlabeled cells. Nuclei in all cells were labeled using 0.75 µM SiR-DNA (Spirochrome, SC007). 24 h after seeding, control samples were left untreated and EMT was induced in the EMT samples by addition of 5 ng/ml HGF (R&D Systems, #294-HG)^33,64,65^.

#### Time-lapse imaging using spinning disk confocal microscopy

Live-cell imaging was performed at 37 °C in CO_2_-independent Leibovitz’s L-15 medium, no phenol red (Thermo Fisher, # 21083027), supplemented with 10% FBS and 1% penicillin-streptomycin. Samples were grown on hydrogels in 2-well chambered coverglasses. Time-lapses were acquired using a 30x silicon oil immersion objective (UPLSAPO30XS, N.A. 1.05, Olympus) on a spinning disk imaging system with a Perkin Elmer MLS1 base laser engine, CSU-X1 Spinning Disk head, Olympus IX81 stand, equipped with a Hamamatsu Flash 4.0v2 sCMOS camera and controlled by Volocity software. Acquired z-stacks had a thickness of 60 µm and a step size of 0.75 µm, the time interval was 10 minutes over a total duration of 10 to 22 h.

### Chemical perturbations

The Rac1 inhibitor NSC23766 (Tocris, #2161) was dissolved and stored as stock solution (10 mM) in DMSO at −20°C. Methyl-β-cyclodextrin (MbCD, Merck, #C4555) was dissolved in cell culture medium and always prepared immediately before experiments. For multi-position live-cell imaging in perturbation experiments, imaging positions were identified and added to Volocity, the inhibitor solution was then added to the imaging medium at the spinning disk confocal microscope and z-stacks of cells were immediately acquired for 30 minutes.

After that the acquisition was interrupted, 5 ng/ml HGF was added to the EMT samples and acquisition was then resumed for 10 to 22 h. In all treatments, cells were seeded on hydrogels 24 h prior to the experiment. Final concentrations used for the drug treatments were 100 µM Rac1 inhibitor NSC23766 and 1mM MbCD.

### Image analysis and morphospace representation

Time lapse movies were cropped in Fiji^66^.

#### Cell segmentation and tracking in 3D

Semi-automatic segmentation and tracking were performed on 3D time lapses of individual cells labeled with Lifeact-GFP using the Surface tool in Imaris 9.1/10 (Bitplane/Oxford Instruments). The Lifeact-GFP (actin) channel was used for segmentation. The following parameters were used in the Surface Tool: Smoothing surface detail length 1.5 µm, background subtraction sphere diameter 8 µm, autoregressive motion tracking with 0 gap size. The faces and vertices of the triangulated surfaces were exported as *.wrl files.

#### Shape quantification

##### Pre-processing

The *.wrl files of the time-dependent shapes of individual cells were imported in MATLAB (versions R2022b-R2024a) with a custom script. Due to segmentation artefacts resulting from regions within the cell with low signal intensity, a cell could consist of multiple meshes. Hence, at each time point, only the largest connected mesh was kept and smaller connected meshes were removed.

##### Standard shape features

We computed standard shape features using a custom script in MATLAB. To calculate the volumetric center of mass 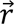_*COM*_ = {*x*_COM_, *y*_COM_, *z*_COM_} of a cell, we used Gauss’s theorem. Cell shape features such as principal axes lengths and directions, cell volume and cell surface area were calculated in a similar manner (Supplemental Information).

##### Spherical harmonics decomposition

We carried out the spherical harmonics decomposition in MATLAB (versions R2022b-R2024a). First, cells were aligned by calculating the eigenvectors and eigenvalues of the covariance matrix of the surface given by

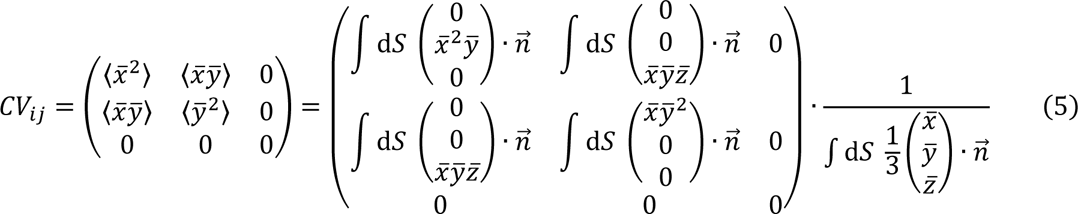

with the centered cell surface points given by *x̅* = *x* − *x*_COM_, *y̅* = *y* − *y*_COM_ and *z̅* = *z* − *z*_COM_. To numerically compute the covariance matrix of a triangulated cell surface mesh, we used Gauss’s theorem (Supplemental Information). The z-direction was chosen perpendicular to the substrate and only the directions in the plane parallel to the substrate were considered, hence we set *CV*_*i*3_ = 0 and *CV*_3*i*_ = 0. Cells were rotated around the z-axis so that the long axis of the cell (corresponding to the eigenvector with the largest eigenvalue) was parallel to the *x*-direction. The initial direction of the cell alignment along the *x*-axis was chosen such that the third moment 〈*x̅*^3^〉 of the mesh was positive. For later time points, the direction of alignment was chosen such that it was pointing in a similar direction as at the previous time point by ensuring that the scalar product of the eigenvectors at the current and previous time point was positive.

After alignment, the MATLAB function *reducepatch()* was used to reduce the number of faces to 1000 per cell, significantly speeding up the following analysis without sacrificing global shape information. For the spherical harmonics decomposition of triangulated shapes, we adapted tools previously developed to study brain structure morphologies and implemented in MATLAB^67,68^ [https://brainimaging.waisman.wisc.edu/~chung/midus/]. First, the surface mesh was mapped onto a unit sphere utilizing an area-preserving surface flattening algorithm^67^ [https://brainimaging.waisman.wisc.edu/~chung/midus/]. This algorithm maps the vertices of the original cell mesh to the surface of a sphere with the aim of having the areas of the faces connecting groups of three vertices being conserved between the original mesh and the spherical mesh. While the algorithm allows for changes in the triangle shape (as long as the area is conserved), the spherical mesh has the same set of edges (connections between two neighboring vertices) as the original mesh, and hence conserves local information of the underlying cell shape. This ensures that similar shapes with different meshes will have similar mappings on the spherical surface. The new vertex positions on the sphere were then rescaled such that they are located on the surface of the unit sphere. The algorithm maps the positions *x*(*θ*, *ϕ*), *y*(*θ*, *ϕ*) and *z*(*θ*, *ϕ*) of the vertices on the unit sphere surface with the polar angle *ϕ* and the azimuthal angle *θ*. These distributions on the surface of the unit sphere were then decomposed into spherical harmonics functions (SPHARMs)^68^ [https://brainimaging.waisman.wisc.edu/~chung/midus/] so that

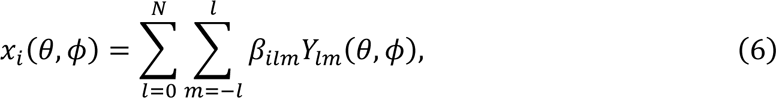

where *Y*_*lm*_(*θ*, *ϕ*) are spherical harmonics functions and *β*_*i,lm*_ are the spherical harmonics coefficients (with *i* = 1,2,3) and which are unique descriptors of cell shape. We also introduced the coordinates *x*_1_, *x*_2_, *x*_3_ which correspond to *x*, *y*, *z*. Here, *N* is the highest degree of spherical harmonics functions considered. The SPHARMs framework also allows for the reconstruction of shapes from the coefficients *β*_*ilm*_with *i* ∈ {1,2,3} (Figure S1A). To verify the quality of the spherical harmonics decomposition, we compared the distances between the triangulated mesh of the original surface and the mesh reconstructed after the spherical harmonics decomposition. For each vertex of an original mesh/reconstructed mesh, we computed the distance from the reconstructed triangulated mesh/original mesh, using the function *point2trimesh()*^69^. We then calculated the maximum of the distances of the vertices and the mean distance of the vertices and meshes of the original and reconstructed meshes.

To identify the minimal degree *N* of spherical harmonic functions (corresponding to 3(*N* + 1)^2^ SPHARMs coefficients, Figure S1B) that represent the shape in the appropriate manner, we performed a spherical harmonics decomposition for values of *N* = 1 … 43 of all cells (without drug treatments) after 0 h and 18 h. For all cells we calculated the maximum error and mean error and remove those cells that have too large errors (maximum error below 2 µm and a mean error below 0.15 µm) for the degree *N* = 30, for which we would usually expect to find a sufficiently small error. For the remaining cells, we calculate the largest mean and maximum error as a function of *N* of all cells. We chose the decomposition degree *N* for which the largest mean error of any cell is below 0.15 µm and the largest maximum error of any cell is below 2 µm [Figures S1C and S1D]. We found that this is fulfilled for *N* = 23.

We then decomposed all cell meshes (for all conditions and times) and only considered those cells with maximum error below 2 µm and a mean error below 0.15 µm for further analysis.

To reconstruct the average shapes of cells as a function of time, we calculated ⟨*β*_*j*,*lm*_⟩_cells_, and reconstructed the resulting cell shape.

#### Dimensionality reduction

For dimensional reduction of the high-dimensional shape features *β*_*ilm*_, we used principal component analysis (PCA). To define the principal components (PCs), we used the shapes features of EMT cells at 0 h and 18 h after HGF addition. We computed their average cell shape features:

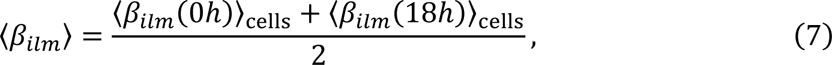

and then defined the centered shape features of the cells as:

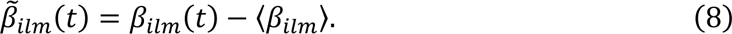

We then used the MATLAB function *pca()* for the PCA of the centered shape features with the observation weights 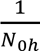 for the cells at 0 h (with *N_0h_* = 128 of cells used for the PCA at 0 h) and the weights 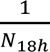 for the cells at 18 h (with *N*_18ℎ_ = 63 of cells used for the PCA at 18 h). The observation weights compensated that at 0 h we have 128 cells that contribute to the PCA and at 18 h we have 63 cells that contributed to the PCA. This provided us with a mapping of the high-dimensional cell shape features into a low-dimensional morphospace. Combining the PC scores derived from the PCA and the shape features 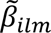(*t*) of all cells, we calculated the PC coefficients (called PC*i* with *i* = 1,2, …) of each individual cell, corresponding to the coordinates in the low-dimensional morphospace In this study, we take the principal components (PCs) to be unitless, such that principal components score has units of [1/µm]. We note that one could instead take the principal component scores of the PCA to be unitless, in which case the PCs would be given in units of [µm]. Using ⟨*β*_*ilm*_⟩ as a reference, we reconstructed cell shapes for arbitrary values of the PC scores, allowing us to reconstruct the cell morphology corresponding to any location in the morphospace, using published MATLAB functions^67,68^ [https://brainimaging.waisman.wisc.edu/~chung/midus/]. To generate bagplots of the cell distributions in the morphospace (Figures 2A, 5B, 5G, S2C, S2D), we employed the LIBRA toolbox^63^.

### Stochastic inference and modelling

#### Numerical inference of deterministic force and diffusion coefficient

Our stochastic inference approach was inspired by previous studies^37^. We assumed that the cell trajectories of the variable PC1 followed overdamped Langevin dynamics

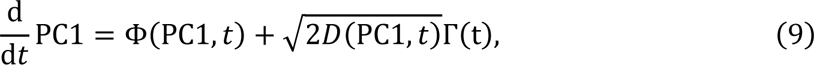

with a deterministic force field Φ(PC1, *t*), the diffusion coefficient *D*(PC1, *t*) and Gaussian white noise Γ(t). Both Φ(PC1, *t*) and *D*(PC1, *t*) can depend on PC1 and time *t*.

We derived the deterministic force field Φ(PC1*t*) from the mean displacement

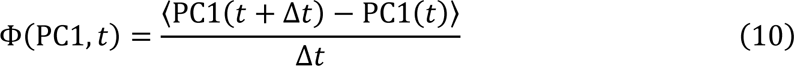

and the diffusion coefficient from the mean squared displacement

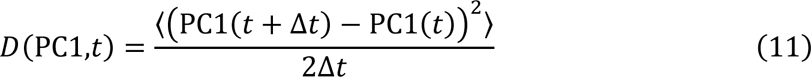

(see Supplemental Information). To numerically infer the deterministic force Φ(PC1,*t*), we considered all trajectories for a given condition (distinguishing HGF and drug treatment, and considering the time interval between 0.5h before EMT induction until 23 h50 min after EMT induction) and considered the ensemble of all their possible displacements (PC1(*t* + Δ*t*) − PC1(*t*))⁄Δ*t*) for the time step Δ*t* = 10 min and fit polynomial surfaces up to the 5^th^ order depending on time *t* and shape PC1(*t*) with the MATLAB function *fit()*. We weighted displacements depending on their time, to make sure that early time points, where we usually have more data, did not overcontribute to the fitting. To identify the best polynomial fit, we computed the degree-of-freedom adjusted coefficient of determination *R*^2^ for each fit (ranging from 0^th^ order to 5^th^ order) and chose the fit with the highest *R*^2^. The same computation was done to derive the diffusion coefficient *D*(PC1, *t*) from the mean squared displacements of the trajectories (PC1(*t* + Δ*t*) − PC1(*t*))^2^ ⁄2Δ*t*. The fitting parameters used in Figures 3C and 3D are provided in the Supplemental Information. When inferring the time-independent behavior of the deterministic force Φ_*m*_(PC1), we only considered polynomials up to the 5^th^ order that depended on PC1. When inferring the shape-independent behavior of the diffusion coefficient *D*_*m*_(t), we only considered polynomials up to the 5^th^ order depending on time *t*.

For the time-independent deterministic force Φ_*m*_(PC1), we derived the corresponding potential using

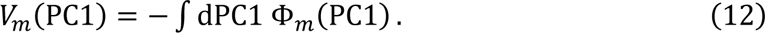

We calculated the 95% prediction intervals for the deterministic forces Φ(PC1, *t*) and Φ_*m*_(PC1) and the diffusion coefficients *D*(PC1, *t*) and *D*_*m*_(t), using the MATLAB function *predint() (*with non-simultaneous bounds for the fitted curve as a function on PC1 and *t*) and estimated the fitting error *e*(PC1, *t*) as difference between the upper and lower prediction bounds. We only considered those regions of PC1 and *t* with

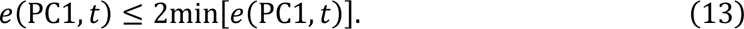

A diffusion coefficient that depends on PC1 (also referred to as multiplicative noise) causes a spurious force

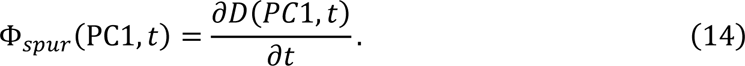

that contributes to the deterministic force. We found that this force contribution is small compared to the deterministic force Φ(PC1, *t*) (Figure S4A).

Changes in PC1 due to deterministic force Φ will translate into changes of the mean squared displacement and hence, can affect the inferred diffusion coefficient. The resulting contribution is given by (Supplemental Information)

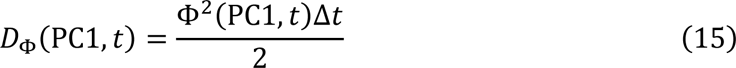

and we found that it has a negligible effect on the inferred diffusion coefficient (Figure S4B).

#### Numerical generation of in silico trajectories of cell shape changes during EMT

To numerically generate stochastic trajectories following Langevin dynamics with the force field Φ(PC1, *t*) and diffusion coefficient *D*(PC1, *t*) extracted from our experimental data as input, we used the numerical scheme^70^

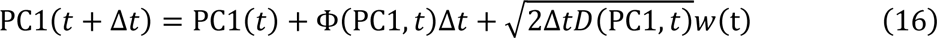

with the starting position PC1(0) and a Gaussian-distributed random variable *w*(*t*) with zero mean and variance 1. We randomly sampled the starting positions of the experimental trajectories as starting position of the simulations and simulated 100 trajectories for each condition with a time step of 0.01 h and saved trajectories of the *ρ*-coordinate with a time interval of 0.17 h.

#### Verification of stochastic inference and *in silico* simulations

To compare numerically generated trajectories with experimentally derived trajectories, for both we simulated several *in silico* trajectories (Figures S3A and S3B) and inferred the deterministic force field and diffusion coefficient from the in silico trajectories. We found good qualitative agreement with the fields inferred from the experimental data (see Figures S3C and S3D). For the inference, we ignored trajectories with shapes PC1 < −30 at 18 h, as these trajectories corresponded to physically invalid cell shapes and were a result from fitting errors in regions with low densities or no cell shapes during the stochastic inference. Moreover, we compared the ensemble behavior for several trajectories ⟨PC1(*t*)⟩ (Figures S3E, S3F) and found good qualitative agreement between the experiment and the simulations.

#### Total diffusion coefficient for first 22 PCs

To calculate the overall shape noise strength of not just PC1, but the first 22 principal components combined (which account for 90% of the total variance of the data, Figure S3G), we defined the high-dimensional morphospace trajectory

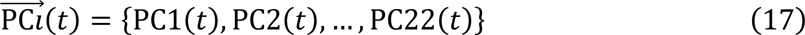

and inferred the diffusion coefficient by fitting polynomial surfaces to the mean squared displacement

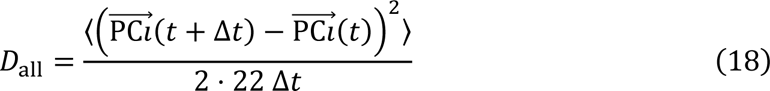

with Δ*t* = 10 min.

#### Time to reach attractor minimum in a simplified mathematical model

We considered a stochastic process expressed by the Langevin equation^36^ of position *ρ* (Figure 4A), which is written as

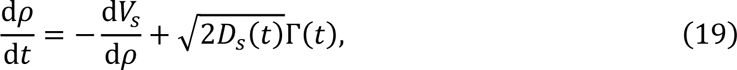

with the initial position *ρ(*0) = *ρ*_ep_ < 0, the quadratic potential *V*_*s*_ = *k*_*s*_*ρ*^2^/2 (corresponding to a potential minimum located at *ρ* = 0), the potential strength *k*_*s*_, and the time-dependent diffusion coefficient (Figure 4B) which is defined as

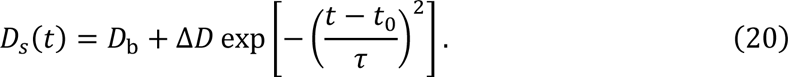

For the diffusion coefficient, we introduced the background noise strength *D*_b_ > 0, the peak height Δ*D* ≥ 0, the peak time *t*_*D*_ and a time *τ* which describes the characteristic time of the peak. The probability of passing the potential minimum located at ρ = 0 at least once after time *t* was then given by

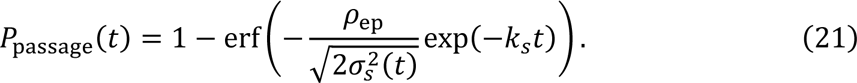

(Figure 4C; Supplemental Information). Here, we introduced the variance

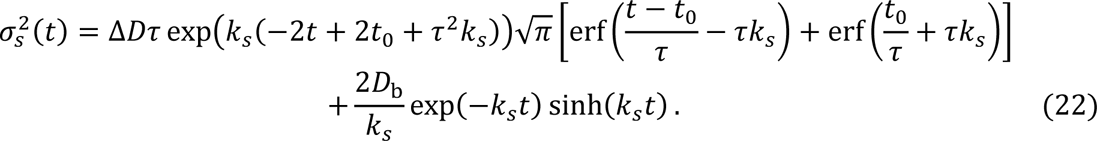

with the error function erf(*x*). The mean first passage time (MFPT) to reach the potential minimum then follows from

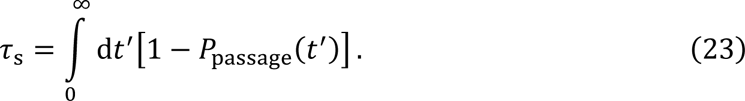

To compute the occupancy of particles in region defined by ρ ∈ [*ρ*_thr_, ∞] (Figure 4A) at time *t* is given by

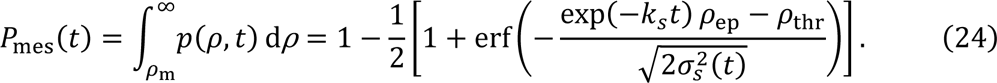

(Figure 4D and Supplemental Information). To calculate the mean first passage time to reach a region defined by ρ ∈ [*ρ*_thr_, ∞] (Figure 4A), we numerically generated trajectories with the starting location *ρ*_ep_ and computed the time when the position of the simulated trajectories is *ρ* ≥ *ρ*_thr_ as a function of the threshold position *ρ*_thr_(Figure 4E).

The parametrization was chosen so that it reflects our experimental observations: *k*_*s*_ = 0.1 h^−1^, *τ* = 5 h and *t*_0_ = 6 h. We consider three cases: constant low noise (*D*_b_ = 3 h^−1^, Δ*D* = 0), constant high noise (*D*_b_ = 16 h^−1^, Δ*D* = 0) and noise peak (*D*_b_ = 3 h^−1^, Δ*D* = 13 h^−1^). We used the initial position *ρ*_ep_ = −15. In Figure 4D, we simulated 500000 trajectories for each case of the diffusion coefficient *D*_*s*_(*t*) and threshold location *ρ*_thr_ with the time step Δ*t* = 0.002 h. To this aim, we employed the numerical scheme provided in Equation 16. To define the mesenchymal region in Figure 4E, we used *ρ*_thr_ = −8 and tested that the presented results are robust when considering other combinations of *ρ*_ep_and *ρ*_thr_.

#### Estimation of average diffusion coefficient

We estimated the average diffusion coefficient (Figure 4I) of an individual cell trajectory of PC 1 of the form {PC1(0), PC1(Δ*t*), PC1(2Δ*t*), PC1(3Δ*t*), PC1(4Δ*t*), … } with the time step Δ*t* = 10 min by calculating the mean squared displacement

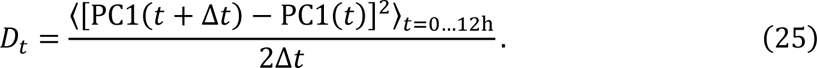

Here, we only considered the first 12 h of the trajectories after induction of EMT via HGF, corresponding to the time until cell spreading in ensemble averaged cell shape (Figure 2D). To define three classes of EMT cell shape trajectories with weak, medium and high noise strength, we considered the distribution of *D*_*t*_ of all EMT cells and considered the 33% and 67% percentiles as the thresholds of the classes.

### Calculations of the cell spreading efficiency parameters

#### Support vector machine classifier

We trained a classifier to assign a mesenchymal shape score to shapes in the PC1-PC2 morphospace. We assumed that at 0 h, all cells had epithelial shapes and at 18 h, all cells had mesenchymal shapes. We then trained a support vector machine classifier in MATLAB with the function *fitcsvm()* with a first order polynomial kernel. For the training, we standardized the predictor data. After training the classifier, we used the *predict()* function in MATLAB to estimate the mesenchymal probability of pairs of PC1 and PC2. We used this function when generating the probability map in Figure 4F and to generate trajectories of mesenchymal shape score of individual cells (e.g., Figure 4G).

#### Estimation of the mean first passage time

We considered 0 h-12 h trajectories (Figures 4H, 4J, 4K, 5E, 5F, 5J, 5K, S5B, S5D and S5E) of the mesenchymal shape score of individual cells, as it corresponds to the time window of cell spreading.

To estimate the probability of cells to reach mesenchymal shapes (Figures 4H, 4J, 5E, 5J, S5B, S5D and S5E), we defined a mesenchymal shape score threshold (between 10% and 100%, Figure 4G) and only considered those trajectories that started with a score below the threshold (assumed to be cells with epithelial shapes). We then measured the percentage of cells that passed this threshold.

To estimate the first passage time (Figures 4K, 5F and 5KI), we considered all trajectories that started with epithelial shapes and passed the threshold and then computed their mean value and standard error.

## QUANTIFICATION AND STATISTICAL ANALYSIS

Statistical tests were conducted using Prism 10.3.0. The number of cells and the statistical tests used are reported in the figure legends. Mann-Whitney test was applied in Fig. 4I.

## SUPPLEMENTAL MOVIE AND DOCUMENT TITLES

Document S1. Model and theoretical calculations.

Movie S1. Times series of cell shape dynamics in control and EMT cells.

Movie S2. Visualization of the shape changes associated with increasing PC1.

Movie S3. Visualization of the shape changes associated with increasing PC2.

Movie S4. Times series of cell shape dynamics in untreated EMT cells and EMT cells with perturbed actin or membrane dynamics.

## Notes

### Competing Interest Statement

The authors have declared no competing interest.

### Summary of Updates

Text, figures and Supplemental files revised.

