## Supplemental Information S1 for "Cell shape noise strength regulates the rate of shape change during EMT-associated cell spreading"

#### 1 Calculation of standard shape features from triangulated cell surfaces

For our morphometric analysis pipeline, we segment cells. This provides us with triangulated cell surface meshes where a cell surface is given by a set of $n$ faces with normal vectors $\vec{n}_i = \{n_{x,i}, n_{y,i}, n_{z,i}\}$ $(i = 1, ..., N)$, face surface areas $S_i$ and face centre of masses $\{x_i, y_i, z_i\}$. The cell volume reads

$$V = \int_{\text{Cell Volume}} \mathrm{d}V = \int_{\text{Cell Surface}} \mathrm{d}S \, \frac{1}{3} \begin{pmatrix} x \\ y \\ z \end{pmatrix} \cdot \vec{n} = \sum_{i=1}^{N} \frac{1}{3} S_i \left( x_i n_{x,i} + y_i n_{y,i} + z_i n_{z,i} \right) \quad \text{(S1)}$$

where we used Gauss's theorem

$$\int_{\text{Cell Volume}} \mathrm{d}V \; \nabla \vec{F} = \int_{\text{Cell Surface}} \mathrm{d}S \; \vec{F} \cdot \vec{n}. \quad \text{(S2)}$$

The centre of mass $\vec{r}_{\text{COM}}$ of the cell surface is computed

$$x_{\text{COM}} = \frac{\int_{\text{Cell Volume}} \mathrm{d}V \; x}{V} = \frac{\int_{\text{Cell Surface}} \mathrm{d}S \begin{pmatrix} 0 \\ xy \\ 0 \end{pmatrix} \cdot \vec{n}}{V} = \frac{1}{V} \sum_{i=1}^{N} S_i x_i y_i n_{y,i}, \quad \text{(S3a)}$$

$$y_{\text{COM}} = \frac{\int_{\text{Cell Volume}} \mathrm{d}V \; y}{V} = \frac{\int_{\text{Cell Surface}} \mathrm{d}S \begin{pmatrix} xy \\ 0 \\ 0 \end{pmatrix} \cdot \vec{n}}{V} = \frac{1}{V} \sum_{i=1}^{N} S_i x_i y_i n_{x,i}, \quad \text{(S3b)}$$

$$z_{\text{COM}} = \frac{\int_{\text{Cell Volume}} \mathrm{d}V \; z}{V} = \frac{\int_{\text{Cell Surface}} \mathrm{d}S \begin{pmatrix} xz \\ 0 \\ 0 \end{pmatrix} \cdot \vec{n}}{V} = \frac{1}{V} \sum_{i=1}^{N} S_i x_i z_i n_{x,i}. \quad \text{(S3c)}$$

To compute the cell covariance matrix, we centred the cells by defining

$$\overline{x}_i = x_i - x_{\mathrm{COM}}, \overline{y}_i = y_i - y_{\mathrm{COM}}, \overline{z}_i = z_i - z_{\mathrm{COM}}. \tag{S4a}$$

and then computed the averages

$$\langle \overline{x}^2 \rangle = \frac{\int_{\text{Cell Volume}} \mathrm{d}V \ \overline{x}^2}{V} = \frac{\int_{\text{Cell Surface}} \mathrm{d}S \begin{pmatrix} 0 \\ \overline{x}^2\overline{y} \\ 0 \end{pmatrix} \cdot \vec{n}}{V} = \frac{1}{V}\sum_{i=1}^{N} S_i \overline{x}_i^2 \overline{y}_i n_{y,i}, \tag{S5a}$$

$$\langle \overline{xy} \rangle = \frac{\int_{\text{Cell Volume}} \mathrm{d}V \ \overline{xy}}{V} = \frac{\int_{\text{Cell Surface}} \mathrm{d}S \begin{pmatrix} 0 \\ 0 \\ \overline{xyz} \end{pmatrix} \cdot \vec{n}}{V} = \frac{1}{V}\sum_{i=1}^{N} S_i \overline{x}_i \overline{y}_i \overline{z}_i n_{z,i}, \tag{S5b}$$

$$\langle \overline{y}^2 \rangle = \frac{\int_{\text{Cell Volume}} \mathrm{d}V \ \overline{y}^2}{V} = \frac{\int_{\text{Cell Surface}} \mathrm{d}S \begin{pmatrix} \overline{xy}^2 \\ 0 \\ 0 \end{pmatrix} \cdot \vec{n}}{V} = \frac{1}{V}\sum_{i=1}^{N} S_i \overline{x}_i \overline{y}_i^2 n_{x,i}. \tag{S5c}$$

The covariance matrix can then be written as

$$\mathrm{CV}_{ij} = \begin{pmatrix} \langle \overline{x}^2 \rangle & \langle \overline{xy} \rangle & 0 \\ \langle \overline{xy} \rangle & \langle \overline{y}^2 \rangle & 0 \\ 0 & 0 & 0 \end{pmatrix}. \tag{S6}$$

As we only considered the orientation of the cell in the $x-y-$plane, we set $\mathrm{CV}_{3j} = 0$ and $\mathrm{CV}_{i3} = 0$. We then computed the eigenvectors of the covariance matrix to define the principal directions. The direction corresponding to the largest eigenvalue was chosen as the new $x$-axis. To define in which direction the cell is pointing, we ensured that the third moment of the aligned coordinates $\{\tilde{x}_i, \tilde{y}_i, \tilde{z}_i\}$ along the $\tilde{x}$-axis is positive:

$$\langle \tilde{x}^3 \rangle = \frac{\int_{\text{Cell Volume}} \mathrm{d}V \ \tilde{x}^3}{V} = \frac{\int_{\text{Cell Surface}} \mathrm{d}S \begin{pmatrix} 0 \\ \tilde{x}^3\tilde{y} \\ 0 \end{pmatrix} \cdot \vec{n}}{V} = \frac{1}{V}\sum_{i=1}^{N} S_i \tilde{x}_i^3 \tilde{y}_i n_{y,i} > 0. \tag{S7}$$

### 2 Stochastic Inference

#### 2.1 Deriving the deterministic force and diffusion coefficient for overdamped Langevin dynamics

We consider the case where the stochastic dynamics of cell shape trajectories. We introduce the variable $\rho = \mathrm{PC1}$ for simplicity. It follows an overdamped Langevin equation

$$\frac{\mathrm{d}}{\mathrm{d}t}\rho = \Phi(\rho, t) + \sqrt{2D_s(\rho, t)}\Gamma(t) \tag{S8}$$

written in the Itô convention [1] with the deterministic force field $\Phi(\rho, t)$, the diffusion coefficient $D(\rho, t)$, the time $t$ and the Gaussian white noise $\Gamma(t)$ satisfying

$$\langle \Gamma(t) \rangle = 0, \tag{S9a}$$

$$\langle \Gamma(t)\Gamma(t') \rangle = \delta(t - t'), \tag{S9b}$$

where $\langle ... \rangle$ denotes averages over multiple realisations. To derive both $\Phi(\rho, t)$ and $D(\rho, t)$ from measurements of $\rho(t)$, we consider the displacement after a small time step $\Delta t$, given by the expansion [2]

$$\rho(t + \Delta t) - \rho(t) \approx \Phi(\rho, t)\Delta t + \sqrt{2D_s(\rho, t)\Delta t}w(t) \tag{S10}$$

where we only consider the lowest order terms of $\Delta t$ in the contributions of $\Phi(\rho, t)$ and $D_s(\rho, t)$, and where we introduced the independent Gaussian-distributed random variables $w(t)$ with mean 0 and variance 1. The mean displacement is given by

$$\langle \rho(t + \Delta t) - \rho(t) \rangle = \Phi(\rho, t)\Delta t \tag{S11}$$

which leads to the deterministic force

$$\Phi(\rho, t) = \frac{\langle \rho(t + \Delta t) - \rho(t) \rangle}{\Delta t}. \tag{S12}$$

To derive the diffusion coefficient, we calculate the mean squared displacement

$$\langle [\rho(t + \Delta t) - \rho(t)]^2 \rangle = (\Phi(\rho, t)\Delta t)^2 + 2\Delta t D_s(\rho, t). \tag{S13}$$

Then, the diffusion coefficient can be calculated from

$$D_s(\rho, t) = \frac{\langle [\rho(t + \Delta t) - \rho(t)]^2 \rangle}{2\Delta t} - \frac{\Phi^2(\rho, t)\Delta t.}{2} \tag{S14}$$

For small $\Delta t$, we only consider the lowest order term including $\Delta t$, leading to

$$D_s(\rho,t) \approx \frac{\langle[\rho(t+\Delta t)-\rho(t)]^2\rangle}{2\Delta t}. \tag{S15}$$

where we ignored the higher-order contribution $-\Phi^2(\rho,t)\Delta t/2$ caused by the deterministic force field $\Phi(\rho,t)$.

#### 2.2 Spurious Force

A diffusion coefficient $D(\rho)$ that depends on $\rho$ causes multiplicative noise that can cause a spurious force contribution to the deterministic force $\Phi(\rho)$. To illustrate this behaviour, we consider a Langevin process

$$\frac{\mathrm{d}}{\mathrm{d}t}\rho = \Phi(\rho) + \sqrt{2D_s(\rho)}\Gamma(t) \tag{S16}$$

with

$$\langle\Gamma(t)\rangle = 0, \tag{S17a}$$
$$\langle\Gamma(t)\Gamma(t')\rangle = \delta(t-t'). \tag{S17b}$$

The corresponding Fokker Planck equation, describing the dynamics of the probability density function $p(\rho)$ of $\rho$, reads [2]

$$\frac{\mathrm{d}}{\mathrm{d}t}p(\rho) = -\frac{\mathrm{d}}{\mathrm{d}\rho}\left(\Phi(\rho)p(\rho)\right) + \frac{\mathrm{d}^2}{\mathrm{d}\rho^2}\left(D_s(\rho)p(\rho)\right). \tag{S18}$$

Its steady-state solution $p_s(\rho)$, resulting from

$$\frac{\mathrm{d}}{\mathrm{d}t}p(\rho) = 0 \tag{S19}$$

then reads

$$p_s(\rho) = C\exp\left(\int_{-\infty}^{\rho}\mathrm{d}\rho'\,\frac{\Phi(\rho') - \frac{\mathrm{d}}{\mathrm{d}\rho'}D_s(\rho')}{D_s(\rho')}\right) \tag{S20}$$

with a normalisaton constant $C$. This distribution will have a peak at location $\rho_p$ when

$$\frac{\mathrm{d}}{\mathrm{d}\rho}p_s(\rho)|_{\rho=\rho_p} = 0. \tag{S21}$$

By calculating

$$\frac{\mathrm{d}}{\mathrm{d}\rho}p_s(\rho) = C\exp\left(\int_{-\infty}^{\rho}\mathrm{d}\rho'\,\frac{\Phi(\rho') - \frac{\mathrm{d}}{\mathrm{d}\rho'}D_s(\rho')}{D_s(\rho')}\right)\frac{\Phi(\rho) - \frac{\mathrm{d}}{\mathrm{d}\rho}D_s(\rho)}{D_s(\rho)} \tag{S22}$$

we find that Equation (S21) is fulfilled when

$$\Phi(\rho_p) - \frac{\mathrm{d}}{\mathrm{d}\rho}D_s(\rho)|_{\rho=\rho_p} = 0 \tag{S23}$$

Here, we introduced the so-called spurious force

$$F_{\text{spurious}}(\rho) = \frac{\mathrm{d}}{\mathrm{d}\rho} D_s(\rho). \tag{S24}$$

A steady state peak in the distribution $p_s(\rho)$ of position $\rho$ needs to be derived from $\Phi(\rho) - F_{\text{spurious}}(\rho)$ and not just from $\Phi(\rho)$. In Figure S4A, we show that $\Phi(\rho) - F_{\text{spurious}}(\rho) \approx \Phi(\rho)$ and thus, the spurious contribution is small.

### 3 Fitting coefficients of the stochastic inference

Below, we provide the fitting parameters of the force and diffusion field in Fig. 3.

#### 3.1 Deterministic force field

For the fitting of the deterministic force field, we use a polynomial of the form

$$\Phi(\rho, t) = \sum_i \sum_j \alpha_{i,j} \rho^i t^j \tag{S25}$$

with the coefficients $\alpha_{i,j}$. The coefficients derived from the stochastic inference (Figure 3C) for the control experiment are given by

| Coefficient | Value | Lower Bound | Upper Bound |
|---|---|---|---|
| $\alpha_{0,0}$ [1/h] | -0.5142 | -1.476 | 0.448 |
| $\alpha_{1,0}$ [1/h] | -0.1816 | -0.3553 | -0.007908 |
| $\alpha_{0,1}$ [1/h$^2$] | 0.8408 | 0.355 | 1.327 |
| $\alpha_{2,0}$ [1/h] | -0.01223 | -0.02895 | 0.004488 |
| $\alpha_{1,1}$ [1/h$^2$] | 0.08097 | 0.009284 | 0.1526 |
| $\alpha_{0,2}$ [1/h$^3$] | -0.1687 | -0.2473 | -0.09006 |
| $\alpha_{3,0}$ [1/h] | 3.032e-05 | -0.001071 | 0.001132 |
| $\alpha_{2,1}$ [1/h$^2$] | 0.004565 | 9.479e-05 | 0.009035 |
| $\alpha_{1,2}$ [1/h$^3$] | -0.01506 | -0.02558 | -0.004541 |
| $\alpha_{0,3}$ [1/h$^4$] | 0.01127 | 0.00644 | 0.0161 |
| $\alpha_{4,0}$ [1/h] | -2.231e-05 | -5.823e-05 | 1.361e-05 |
| $\alpha_{3,1}$ [1/h$^2$] | 0.0001606 | -7.589e-06 | 0.0003289 |
| $\alpha_{2,2}$ [1/h$^3$] | -0.0004072 | -0.0007991 | -1.532e-05 |
| $\alpha_{1,3}$ [1/h$^4$] | 0.0009684 | 0.0003546 | 0.001582 |
| $\alpha_{0,4}$ [1/h$^5$] | -0.0002444 | -0.0003436 | -0.0001451 |
| $\alpha_{5,0}$ [1/h] | -2.695e-06 | -4.006e-06 | -1.384e-06 |
| $\alpha_{4,1}$ [1/h$^2$] | 1.191e-06 | -1.563e-06 | 3.945e-06 |
| $\alpha_{3,2}$ [1/h$^3$] | -5.816e-06 | -1.254e-05 | 9.105e-07 |
| $\alpha_{2,3}$ [1/h$^4$] | 1.101e-05 | 9.058e-07 | 2.112e-05 |
| $\alpha_{1,4}$ [1/h$^5$] | -2.03e-05 | -3.253e-05 | -8.066e-06 |

Table 1: Fitting coefficients $\alpha_{i,j}$ with 95% confidence bounds for the control experiment.

For the EMT experiment (Figure 3C), it is given by

| Coefficient | Value | Lower Bound | Upper Bound |
|---|---|---|---|
| $\alpha_{0,0}$ [1/h] | 1.503 | 0.436 | 2.57 |
| $\alpha_{1,0}$ [1/h] | 0.1165 | -0.02514 | 0.2582 |
| $\alpha_{0,1}$ [1/h$^2$] | -0.03837 | -0.5865 | 0.5097 |
| $\alpha_{2,0}$ [1/h] | -0.003008 | -0.01395 | 0.007932 |
| $\alpha_{1,1}$ [1/h$^2$] | -0.0673 | -0.1262 | -0.008443 |
| $\alpha_{0,2}$ [1/h$^3$] | 0.04637 | -0.04654 | 0.1393 |
| $\alpha_{3,0}$ [1/h] | -0.0005881 | -0.001093 | -8.305e-05 |
| $\alpha_{2,1}$ [1/h$^2$] | 0.0003481 | -0.002308 | 0.003004 |
| $\alpha_{1,2}$ [1/h$^3$] | 0.003833 | -0.005652 | 0.01332 |
| $\alpha_{0,3}$ [1/h$^4$] | -0.005721 | -0.01166 | 0.0002142 |
| $\alpha_{4,0}$ [1/h] | 9.229e-06 | -1.553e-05 | 3.399e-05 |
| $\alpha_{3,1}$ [1/h$^2$] | 0.0001409 | 4.56e-05 | 0.0002362 |
| $\alpha_{2,2}$ [1/h$^3$] | -0.0001589 | -0.0004245 | 0.0001068 |
| $\alpha_{1,3}$ [1/h$^4$] | 0.0001676 | -0.0004414 | 0.0007765 |
| $\alpha_{0,4}$ [1/h$^5$] | 0.0001673 | 4.156e-05 | 0.0002931 |
| $\alpha_{5,0}$ [1/h] | -1.011e-07 | -5.527e-07 | 3.505e-07 |
| $\alpha_{4,1}$ [1/h$^2$] | -8.422e-07 | -1.916e-06 | 2.317e-07 |
| $\alpha_{3,2}$ [1/h$^3$] | -4.689e-06 | -8.428e-06 | -9.51e-07 |
| $\alpha_{2,3}$ [1/h$^4$] | 7.178e-06 | -8.63e-07 | 1.522e-05 |
| $\alpha_{1,4}$ [1/h$^5$] | -1.061e-05 | -2.375e-05 | 2.534e-06 |

Table 2: Fitting coefficients $\alpha_{i,j}$ with 95% confidence bounds for the EMT experiment.

#### 3.2 Diffusion field

For the fitting of the diffusion coefficient, we use a polynomial of the form

$$D(\rho, t) = \sum_i \sum_j \beta_{i,j} \rho^i t^j \tag{S26}$$

with the coefficients $\beta_{i,j}$. The coefficients derived from the stochastic inference (Figure 3D) for the control experiment are given by

| Coefficient | Value | Lower Bound | Upper Bound |
|---|---|---|---|
| $\beta_{0,0}$ [1/h] | 5.295 | 4.08 | 6.51 |
| $\beta_{1,0}$ [1/h] | 0.143 | -0.06747 | 0.3535 |
| $\beta_{0,1}$ [1/h$^2$] | 1.443 | 0.6429 | 2.242 |
| $\beta_{2,0}$ [1/h] | -0.0224 | -0.04243 | -0.002375 |
| $\beta_{1,1}$ [1/h$^2$] | 0.03547 | -0.05338 | 0.1243 |
| $\beta_{0,2}$ [1/h$^3$] | -0.3922 | -0.5785 | -0.206 |
| $\beta_{3,0}$ [1/h] | 0.0016 | 0.0002855 | 0.002914 |
| $\beta_{2,1}$ [1/h$^2$] | -0.008089 | -0.01346 | -0.002718 |
| $\beta_{1,2}$ [1/h$^3$] | -0.004797 | -0.01791 | 0.008314 |
| $\beta_{0,3}$ [1/h$^4$] | 0.03441 | 0.01577 | 0.05305 |
| $\beta_{4,0}$ [1/h] | 0.0002676 | 0.000225 | 0.0003102 |
| $\beta_{3,1}$ [1/h$^2$] | -0.0006032 | -0.0008042 | -0.0004023 |
| $\beta_{2,2}$ [1/h$^3$] | 0.001002 | 0.0005319 | 0.001472 |
| $\beta_{1,3}$ [1/h$^4$] | 0.000339 | -0.0004248 | 0.001103 |
| $\beta_{0,4}$ [1/h$^5$] | -0.001319 | -0.002152 | -0.0004868 |
| $\beta_{5,0}$ [1/h] | 6.572e-06 | 5.017e-06 | 8.127e-06 |
| $\beta_{4,1}$ [1/h$^2$] | -1.738e-05 | -2.065e-05 | -1.412e-05 |
| $\beta_{3,2}$ [1/h$^3$] | 1.925e-05 | 1.122e-05 | 2.729e-05 |
| $\beta_{2,3}$ [1/h$^4$] | -2.506e-05 | -3.717e-05 | -1.295e-05 |
| $\beta_{1,4}$ [1/h$^5$] | -8.023e-06 | -2.316e-05 | 7.117e-06 |
| $\beta_{0,5}$ [1/h$^6$] | 1.893e-05 | 5.294e-06 | 3.257e-05 |

Table 3: Fitting coefficients $\beta_{i,j}$ with 95% confidence bounds for the control experiment.

For the EMT experiment (Figure 3D), it is given by

| Coefficient | Value | Lower Bound | Upper Bound |
|---|---|---|---|
| $\beta_{0,0}$ [1/h] | 7.764 | 5.372 | 10.16 |
| $\beta_{1,0}$ [1/h] | 0.1395 | -0.178 | 0.457 |
| $\beta_{0,1}$ [1/h$^2$] | 4.702 | 3.473 | 5.93 |
| $\beta_{2,0}$ [1/h] | -0.007326 | -0.03185 | 0.01719 |
| $\beta_{1,1}$ [1/h$^2$] | 0.3344 | 0.2025 | 0.4663 |
| $\beta_{0,2}$ [1/h$^3$] | -0.8442 | -1.052 | -0.6359 |
| $\beta_{3,0}$ [1/h] | 0.0003908 | -0.0007414 | 0.001523 |
| $\beta_{2,1}$ [1/h$^2$] | 0.00335 | -0.002602 | 0.009303 |
| $\beta_{1,2}$ [1/h$^3$] | -0.0617 | -0.08296 | -0.04044 |
| $\beta_{0,3}$ [1/h$^4$] | 0.04547 | 0.03217 | 0.05878 |
| $\beta_{4,0}$ [1/h] | -3.23e-05 | -8.779e-05 | 2.319e-05 |
| $\beta_{3,1}$ [1/h$^2$] | -2.552e-06 | -0.0002161 | 0.000211 |
| $\beta_{2,2}$ [1/h$^3$] | -0.0002763 | -0.0008717 | 0.0003191 |
| $\beta_{1,3}$ [1/h$^4$] | 0.003675 | 0.00231 | 0.00504 |
| $\beta_{0,4}$ [1/h$^5$] | -0.000783 | -0.001065 | -0.0005011 |
| $\beta_{5,0}$ [1/h] | -1.299e-07 | -1.142e-06 | 8.823e-07 |
| $\beta_{4,1}$ [1/h$^2$] | 2.392e-06 | -1.524e-08 | 4.799e-06 |
| $\beta_{3,2}$ [1/h$^3$] | -1.384e-06 | -9.763e-06 | 6.996e-06 |
| $\beta_{2,3}$ [1/h$^4$] | 6.416e-06 | -1.161e-05 | 2.444e-05 |
| $\beta_{1,4}$ [1/h$^5$] | -7.109e-05 | -0.0001005 | -4.163e-05 |

Table 4: Fitting coefficients $\beta_{i,j}$ with 95% confidence bounds for the EMT experiment.

#### 4 Minimal stochastic model of noise peak

We consider a stochastic process expressed by the Langevin equation

$$\frac{\mathrm{d}}{\mathrm{d}t}\rho = -\frac{\mathrm{d}}{\mathrm{d}\rho}V_s(\rho) + \sqrt{2D_s(\rho)}\Gamma(t) \tag{S27}$$

written in the Itô convention. Here, we introduced $\rho$ as cell shape descriptor, time $t$, the potential $V_s$, the diffusion coefficient $D_s$ and the Gaussian white noise

$$\langle\Gamma(t)\rangle = 0, \tag{S28a}$$
$$\langle\Gamma(t)\Gamma(t')\rangle = \delta(t-t'). \tag{S28b}$$

The initial position reads

$$\rho(0) = \rho_{\mathrm{ep}} < 0. \tag{S29}$$

For this model, we consider the potential

$$V_s(\rho) = \frac{1}{2}k_s\rho^2, \tag{S30}$$

with the time-dependent diffusion coefficient

$$D_s(t) = D_{\mathrm{b}} + \Delta D \exp\left[-\left(\frac{t-t_0}{\tau}\right)^2\right], \tag{S31}$$

with the diffusion background $D_{\mathrm{b}} > 0$, the peak amplitude $\Delta D \geq 0$, the peak time $t_0$ and the peak width $\tau > 0$. Together, this leads to the Langevin equation

$$\frac{\mathrm{d}}{\mathrm{d}t}\rho = -k_s\rho + \sqrt{2\left(D_{\mathrm{b}} + \Delta D \exp\left[-\left(\frac{t-t_0}{\tau}\right)^2\right]\right)}\Gamma(t). \tag{S32}$$

##### 4.1 Solution of the Fokker-Planck equation

The corresponding Fokker-Planck equation for the probability density $p(\rho, t)$ of $\rho$ reads

$$\frac{\mathrm{d}}{\mathrm{d}t}p(\rho,t) = k_s\frac{\mathrm{d}}{\mathrm{d}\rho}\left(\rho p(\rho,t)\right) + \left(D_{\mathrm{b}} + \Delta D \exp\left[-\left(\frac{t-t_0}{\tau}\right)^2\right]\right)\frac{\mathrm{d}^2}{\mathrm{d}\rho^2}p(\rho,t), \tag{S33a}$$

$$p(\rho,0) = \delta(\rho-\rho_{\mathrm{ep}}). \tag{S33b}$$

To solve this partial differential equation, we use the Fourier transform with respect to $\rho$

$$p(\rho, t) = \frac{1}{2\pi} \int_{-\infty}^{\infty} \mathrm{d}k \ \exp(ik\rho)\tilde{p}(k, t), \tag{S34a}$$

$$\tilde{p}(k, t) = \int_{-\infty}^{\infty} \mathrm{d}\rho \ \exp(-ik\rho)p(\rho, t), \tag{S34b}$$

which leads to the rewritten differential equation in Fourier space

$$\frac{\mathrm{d}}{\mathrm{d}t}\tilde{p}(k, t) = -k_s k \frac{\mathrm{d}}{\mathrm{d}k}\tilde{p}(k, t) - \left( D_\mathrm{b} + \Delta D \exp\left[ -\left( \frac{t - t_0}{\tau} \right)^2 \right] \right) k^2 \tilde{p}(k, t), \tag{S35a}$$

$$\tilde{p}(k, 0) = \exp\left(-ik\rho_\mathrm{ep}\right). \tag{S35b}$$

The solution in Fourier space is given by

$$\tilde{p}(k, t) = \exp\left[ -\frac{1}{2} k \left( k\sigma_s^2(t) + 2i\rho_\mathrm{ep} \exp\left(-k_s t\right) \right) \right] \tag{S36}$$

with

$$\sigma_s^2(t) = \Delta D \tau \exp\left[ k_s \left( -2t + 2t_0 + \tau^2 k_s \right) \right] \sqrt{\pi} \left[ \mathrm{erf}\left( \frac{t - t_0}{\tau} - \tau k_s \right) + \mathrm{erf}\left( \frac{t_0}{\tau} + \tau k_s \right) \right] + \frac{D_\mathrm{b}}{k_s} \left[ 1 - \exp\left(-2k_s t\right) \right] \tag{S37}$$

and where we introduced the error function $\mathrm{erf}(x)$. The first term originates from the noise peak $\Delta D$, while the second term is linked to the constant diffusion background $D_\mathrm{b}$. For $t \to \infty$ it converges to

$$\sigma_s^2(t \to \infty) = \frac{D_\mathrm{b}}{k_s}. \tag{S38}$$

A Fourier transform then gives the solution

$$p(\rho, t) = \frac{1}{\sqrt{2\pi\sigma_s^2(t)}} \exp\left[ -\left( \frac{\rho - \rho_\mathrm{ep} \exp\left(-k_s t\right)}{\sqrt{2\sigma_s^2(t)}} \right)^2 \right] \tag{S39}$$

which corresponds to a normal distribution with the mean value

$$\langle \rho(t) \rangle = \rho_\mathrm{ep} \exp\left(-k_s t\right) \tag{S40}$$

and the variance

$$\langle \rho^2(t) \rangle - \langle \rho(t) \rangle^2 = \sigma_s^2(t). \tag{S41}$$

#### 4.2 Probability of reaching the potential minimum and mean first passage time

To calculate the percentage $P_{\text{passage}}$ that a particle with position $\rho(t)$ has at least once passed the potential minimum (located at $\rho = 0$) within the time $t$, we first need to solve the differential equation in Equation (S33) with an absorbing wall at the minimum, corresponding to the boundary condition

$$p(0, t) = 0. \tag{S42}$$

To solve this equation, we utilised the method of images [3], which introduces

$$p_{\text{image}}(\rho, t) = p(\rho, t) - p(-\rho, t). \tag{S43}$$

This equation fulfils Equation (S33), Equation (S42) and $p_{\text{image}}(\rho, t) \geq 0$. The survival probability, corresponding to the probability that a stochastic trajectory $\rho(t)$ has not passed the minimum at time $t$, is given by

$$\begin{aligned} P_{\text{survival}}(t) &= \int_{-\infty}^{0} \mathrm{d}\rho \; p_{\text{image}}(\rho, t) \\ &= \operatorname{erf}\left[-\frac{\rho_{\text{ep}}}{\sqrt{2\sigma_s^2(t)}} \exp\left(-k_s t\right)\right] \end{aligned} \tag{S44}$$

with $\rho_{\text{ep}} < 0$. This leads to

$$\begin{aligned} P_{\text{passage}}(t) &= 1 - P_{\text{survival}} \\ &= 1 - \operatorname{erf}\left[-\frac{\rho_{\text{ep}}}{\sqrt{2\sigma_s^2(t)}} \exp\left(-k_s t\right)\right]. \end{aligned} \tag{S45}$$

#### 4.3 Occupancy of a target region

We next consider the occupancy of a particle with position $\rho(t)$ that started at $\rho(0) = \rho_{\text{ep}} < 0$ and being located within a target region $[\rho_{\text{thr}}, \infty]$ after time $t$ with $\rho_{\text{ep}} < \rho_{\text{thr}} < 0$. Using Eq. (S39), the occupancy of $\rho$ in the target region is

$$\begin{aligned} P_{\text{mes}}(t) &= \int_{\rho_{\text{thr}}}^{\infty} \mathrm{d}\rho \; p(\rho, t) \\ &= 1 - \frac{1}{2}\left(1 + \operatorname{erf}\left[-\frac{\rho_{\text{ep}} \exp(-k_s t) - \rho_{\text{thr}}}{\sqrt{2\sigma_s^2(t)}}\right]\right). \end{aligned} \tag{S46}$$

For $t \to \infty$, this converges to

$$P_{\text{mes}}(t \to \infty) = 1 - \frac{1}{2}\left(1 + \operatorname{erf}\left[\sqrt{\frac{k_s}{2D_{\text{b}}}} \rho_{\text{thr}}\right]\right). \tag{S47}$$
